# Endothelial gain of PIEZO/TRPV mechanosensors drives cavernous malformations

**DOI:** 10.64898/2026.10.01.756057

**Authors:** J de Jong, C. Pasquier, T. Manicom Ramsamy, A. Ahmed, A. Onisiforou, C. Roedel, F. Senger, A.P. Bouin, S. Leverrier-Penna, S. E. Rey, S. Vorsselmans, C. Poveda Vicedo, E. Bryant, A. Shapeti, N. Kroenke, F. Feuerhake, C. Hartmann, I. Petrakakis, C. Oddou, D. Linhares, R. Cardinaels, E. Planus, S. Rocha, A. Samii, H. Bertalanffy, O. Destaing, S. Abdelilah-Seyfried, A. Penna, S. Kar, H. Van Oosterwyck, E. Faurobert

## Abstract

Vascular disease often begins as a failure to read mechanical force. Cerebral cavernous malformations (CCMs) are leaky, brain-invading capillary clusters that cause stroke and seizures and arise from endothelial loss of the CCM complex. They form exclusively in low-flow vessels but existing models cannot explain this selectivity. We previously established that CCM-mutant endothelium inverts its adhesive balance, weakening cell-cell junctions while building matrix-degrading adhesions. The mechanosensors imposing this switch are unknown. Here we show that the mechanosensitive Ca²⁺ channels PIEZO1, TRPV2 and TRPV4 are overexpressed in the endothelium of CCM lesions in a 42-patient cohort, and that lesion volume scales with their expression level. Mechanistically, PIEZO1 partners with TRPV4 to destabilise cell–cell junctions and with TRPV2 to reinforce cell–matrix adhesions; silencing TRPV4 restores junctional integrity and abolishes KLF4 induction, whereas silencing TRPV2 or inhibition with tranilast abolishes matrix degradation and invasion. In vivo, trpv4 knockdown or inhibition with GSK2798745 restores atrioventricular canal morphogenesis in ccm2-mutant zebrafish. Gain of mechanosensory channel function is thus a causal step in CCM pathogenesis, upstream of the KLF program. Combined therapeutic targeting of TRPV2 and TRPV4 with inhibitors validated in humans offers a route to therapy.

## Introduction

Vascular disorders often stem from impaired mechanosensing or mechanoresponse to shear stress, cyclic strain, or wall stiffness signals^1^. Cerebral cavernous malformations (CCMs) is one such disease where defective cell mechanics combines with genetic defects to give rise to clusters of abnormally dilated, leaky capillaries that invade the brain and are prone to leakage, rupture and hemorrhage. Currently, the only treatment option is surgical excision of symptomatic lesions, but this carries substantial risks, particularly for CCMs located in deep brain regions or the brainstem².

CCMs arise from biallelic loss-of-function mutations in any of the three *CCM* genes (*CCM1*, *CCM2*, or *CCM3*)^2,3^, which synergize with low shear stress to drive lesion formation exclusively in low-flow capillaries^4–6^. These lesions exhibit a dynamic and progressive natural history, marked by onset, expansion, and recurrent hemorrhages over the patient’s lifespan^2,7^. The prevailing model posits that CCMs initiate from clonal expansion of a homozygous mutant endothelial cell (EC), forming an immature lesion with focal dilation of a single cavern^8,9^. Mutant ECs then invade the brain parenchyma during a phase of aberrant invasive angiogenesis, recruiting wild-type ECs and inflammatory cells to form genetically mosaic, multicavernous lesions^8–10^.

Ultrastructural analyses have revealed that CCM lesions exhibit endothelial cell–cell junction disruptions and an abnormal basal lamina^11^. In vitro, loss of the CCM protein complex disrupts adherens junctions, increases ECM adhesion, and induces aberrant mechanical and proteolytic ECM remodeling^12,13^. These defects are driven by dysregulated mechanical pathways controlling VE-cadherin- and integrin-dependent adhesions, involving the RhoA effectors ROCK1 and ROCK2^14–16^. Increased ROCK-dependent cellular forces and ECM degradation further facilitate angiogenic invasion^13,17^.

A striking and unresolved feature of CCMs is their selective formation in low-flow venous capillaries, while high flow appears protective^4,18^. *In vitro*, the morphological and transcriptomic defects caused by CCM protein loss manifest exclusively under low-flow or static conditions and are mitigated by high flow^5^. The mechanisms underlying this failure of CCM-mutant ECs to adapt to low flow remain unclear. One plausible explanation is a dysregulation of EC ability to sense the mechanical aspects of blood flow and/or ECM stiffness. Mechanosensitive calcium channels (MSCs), such as those from the PIEZO and TRPV (Transient Receptor Potential Vanilloid) families, have emerged as key regulators of adhesion, migration, and invasion through their control of actin cytoskeleton remodeling, focal adhesion dynamics, and ECM-degrading protease secretion^19,20^. Over the past decade, dysregulated activity and/or expression of Ca²⁺-permeable channels has been linked to enhanced migratory and invasive capacities in tumor cells, leading to their emergence as prognostic markers in various cancers^21^. However, their precise roles in maintaining vascular integrity and regulating angiogenesis remain underexplored.

Here, we demonstrate that PIEZO1, TRPV2, and TRPV4 are upregulated in ECs bordering CCM lesions in a cohort of 42 patients. Furthermore, we show that the TRPV2:TRPV4 expression ratio correlates with the size of the lesions located in the brainstem. A possible molecular explanation comes from our demonstration of the roles of these channels in disrupting endothelial integrity and promoting invasive sprouting angiogenesis. Mechanistically, we show that TRPV4 destabilizes intercellular junctions and increases their permeability by remodeling the actin cytoskeleton while TRPV2 promotes ECM adhesion, degradation and invasion. Moreover, we reveal a functional coupling between PIEZO1 and TRPV4 at intercellular junctions, and between PIEZO1 and TRPV2 at cell–ECM adhesion sites. Additionally, TRPV4 overexpression leads to an upregulation of the mechanosensitive transcription factor KLF4 which has been implicated in causing *in vivo* CCM defects in animal models. Consistent with this, downregulation or pharmacological inhibition of Trpv4 rescues morphogenetic events during cardiac development in *ccm2*-mutant zebrafish embryos.

Overall, these findings suggest that combined therapeutic targeting of TRPV2 and TRPV4 with clinically established safe inhibitors could restore endothelial barrier function while suppressing aberrant angiogenesis in CCM patients, thereby reducing lesion burden. Furthermore, the TRPV2:TRPV4 ratio emerges as a promising avenue in the search for biomarkers of disease progression.

## Results

### Mechanosensitive PIEZO1, TRPV2 and TRPV4 channels are overexpressed in endothelial cells of human CCM lesions

While there is increasing evidence supporting the roles of MSCs in maintaining vascular integrity and regulating angiogenesis, their links with vascular malformations remain understudied. To investigate their potential dysregulation in CCM, we first reanalyzed our previously published RNA sequencing data, focusing on differentially expressed protein-coding genes between control and CCM patients samples^22^. Specifically, we examined members of the TRP and PIEZO channel families, as several TRP channels are known to be mechanosensitive, and PIEZO channels are prototypical MSCs. Among the nine TRP channels identified, TRPV2 and TRPV4, two non-selective cation channels that primarily conduct Ca²⁺ ions into cells, showed the highest and most consistent upregulation across CCM patients (Figure 1a). PIEZO1 also showed a tendency toward overexpression, whereas PIEZO2 expression did not display consistent variation between control and CCM patients.

**Figure 1.**
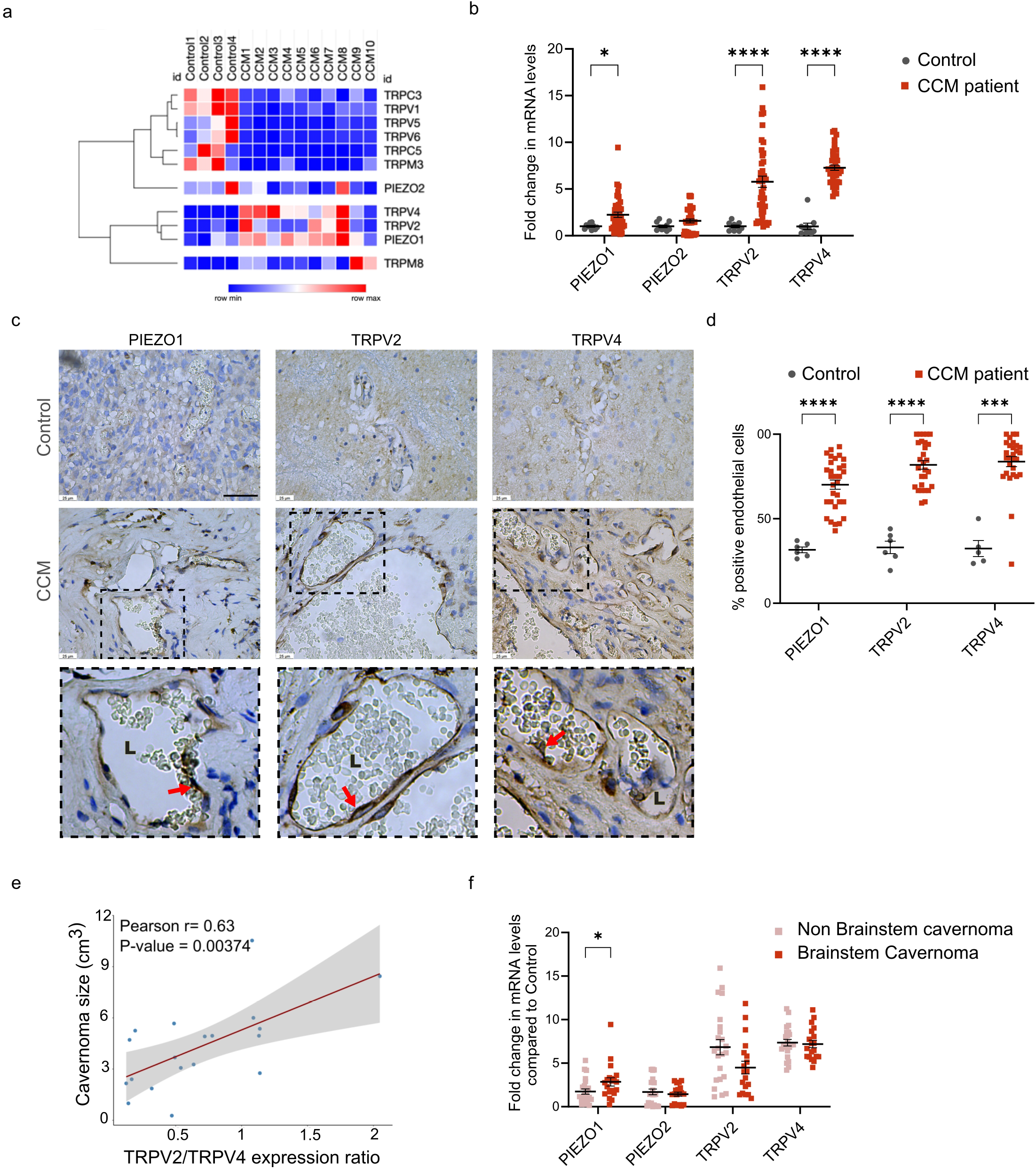
Endothelial PIEZO1, TRPV2, and TRPV4 are overexpressed in human CCM lesions. **(a)** Heatmap of relative mRNA expression (row min to row max) of TRP channel family members (TRPV2, TRPV4, TRPC3, TRPV1, TRPV5, TRPV6, TRPC5, TRPM3, TRPM8) and PIEZO1/PIEZO2 across controls and CCM patient samples, reanalyzed from a previously published differential gene expression dataset. This heatmap was generated using Morpheus software from Broad institute (https://software.broadinstitute.org/morpheus/)^22^. Columns represent individual control and CCM samples. **(b)** Fold change of PIEZO1, PIEZO2, TRPV2 and TRPV4 mRNA levels in CCM lesions (n = 42, red) relative to control brain tissue (n = 10, gray), measured by qRT-PCR and normalized to the endothelial marker CDH5 (VE-cadherin) to account for differences in endothelial cell content between tissues. Each dot represents an individual patient; bars indicate means. Mann Whitney test; *p < 0.05, ****p < 0.0001. Exact p-values are: TRPV2: <0.0001; TRPV4: <0.0001; PIEZO1: 0.0283; PIEZO2: 0.9392 between controls and CCM patients. **(c)** Representative immunohistochemistry for PIEZO1, TRPV2, and TRPV4 in control brain tissue (n=6, top row) and CCM lesion tissue (n=30, middle row), performed with automated staining on CCM paraffin-embedded sections and commercial healthy controls. Bottom row shows magnified views of boxed region; red arrows marking positive endothelial cells. Scale bar, 25 µm. **(d)** Semi-quantitative analysis of PIEZO1, TRPV2, and TRPV4 positive endothelial cells in IHC sections. Each dot represents one section; bars indicate mean ± SEM. Mann-Whitney U test; ***p < 0.001, ****p < 0.0001. Exact p-values are: PIEZO1: 2.49751E-13, TRPV2: 1.61057E-07, TRPV4: 8.15757E-06. **(e)** Positive correlation between TRPV2:TRPV4 mRNA expression ratio and lesion volume (cm³) in patients with brainstem CCM lesions (n = 19). Pearson r = 0.63, p = 0.00374; red line, linear regression; 95% confidence interval. **(f)** PIEZO1, PIEZO2, TRPV2, and TRPV4 fold change in mRNA levels compared to control (qRT-PCR, CDH5-normalized), in non-brainstem (n = 21) versus brainstem (n = 19) lesions. Bars indicate mean ± SEM. Wilcoxon test; *p < 0.05. Exact p-value is p = 0.035 for PIEZO1.

To validate these findings, we analyzed PIEZO1, PIEZO2, TRPV2, and TRPV4 transcript levels in a new and extended cohort of 42 resected CCM lesions (familial and sporadic) and 10 control brain tissues using quantitative PCR (qPCR). Normalization to GAPDH confirmed significant upregulation of PIEZO1, TRPV2 and TRPV4 in CCM samples, but not PIEZO2 (Fig. S1a). To account for global changes in the EC compartment between control and CCM tissues, we normalized the data to CDH5 (VE-cadherin), an endothelial specific marker previously shown to be slightly upregulated in ccm3 KO mice^23^ or in CCM1-3 vessel organoids^24^, so that normalizing to it would only underestimate any increase. Even with this conservative normalization, TRPV2 and TRPV4 remained significantly overexpressed in CCM tissues (Fig. 1b), while PIEZO1 showed a moderate but statistically significant increase. PIEZO2 expression was unchanged. These results suggest a selective upregulation of certain Ca²⁺ channels, particularly members of the TRPV family, in EC cells in the context of the CCM pathology.

To further confirm protein-level upregulation of PIEZO1, TRPV2 and TRPV4, immunohistochemistry was performed using validated antibodies and automated staining on paraffin-embedded sections of 30 CCM lesions concomitantly with 6 control brain tissues. All 3 channels were located in the ECs with stronger staining in ECs bordering the CCM lesions (Fig. 1c). Semi-quantitative analysis using automated threshold revealed a strong increase in the percentage of signal-positive ECs in patient samples compared to control brain tissue (Fig. 1d). These results confirm that PIEZO1, TRPV2 and TRPV4 are upregulated at both mRNA and protein levels in the ECs bordering the CCM lesions.

### TRPV2:TRPV4 ratio correlates with brainstem CCM lesion size

To establish a potential link between specific MSC expression and clinical pathological parameters of CCM disease, we analyzed this cohort of 42 patients with documented lesion volume, location (brainstem vs. non-brainstem), multiplicity, and familial vs. sporadic presentation (Table S1). We observed a moderate but significant positive correlation between TRPV2 levels and CCM lesion size (Pearson *r* = 0.36, *p* = 0.021) (Fig. S1b). Strikingly, the TRPV2:TRPV4 expression ratio showed an even stronger correlation with lesion size in the brainstem (Pearson *r* = 0.63, *p* = 0.00374) (Fig. 1e), a confined region with the highest stiffness in the brain^25^. A positive trend was also observed across all brain regions (Pearson *r* = 0.28, *p* = 0.0682), the lack of significance likely caused by the limited sample size. These data suggest that a higher TRPV2:TRPV4 ratio is associated with increased lesion size, particularly in the brainstem, supporting its potential use as a composite biomarker.

We next compared channel expression between patients with and without brainstem lesions. PIEZO1 expression was significantly elevated in individuals with brainstem involvement (*p* = 0.035) (Fig. 1f), suggesting an association between PIEZO1 upregulation and brainstem-specific CCM localization.

Overall, these analyses suggest that PIEZO1 overexpression may favor CCM initiation in the mechanically challenged brainstem, while TRPV2 and TRPV4 cooperate to drive lesion expansion.

### PIEZO1, TRPV2, and TRPV4 are also upregulated in CCM2-depleted ECs

The intriguing predictive value of these Ca²⁺ channels for CCM lesion burden, prompted to further investigate their role in the endothelial defects associated with the disease. To do so, we employed our well-characterized 2D and 3D *in vitro* models recapitulating EC defects observed in CCM^12–14,17^. These models rely on the use of primary ECs, *i.e*. Human Umbilical Vein Endothelial Cells (HUVEC) and primary Human Brain Microvascular Endothelial Cells (HBMVECs) silenced for CCM2, which is central for the CCM1/2 complex as its loss also leads to the loss of CCM1 protein (KRIT1)^12^.

We first confirmed that PIEZO1, TRPV2, and TRPV4 were upregulated at both the mRNA (Fig. 2a, b) and protein level (Fig. 2c–e) in CCM2-silenced ECs. PIEZO2 was also significantly upregulated in HUVECs but in a more variable manner in HBMVECs. Of note, the lack of commercially available specific antibodies against PIEZO2 prevented us from verifying its expression at the protein level. To assess channel functionality, we measured Ca²⁺ signals in response to specific agonists: Yoda1 for PIEZO1 and GSK1016790A for TRPV4. CCM2-silenced HUVECs exhibited enhanced Ca²⁺ mobilization kinetics through PIEZO1 (Fig. 2f, g) and TRPV4 (Fig. 2i, j), with increased slopes and maximal responses. These effects were abolished or strongly reduced upon additional silencing of TRPV4 or PIEZO1, respectively (Fig. 2f, i), confirming signal specificity. Unfortunately, we could not measure TRPV2-dependent Ca²⁺ signals due to the lack of a commercially available specific pharmacological agonist for the human channel.

**Figure 2:**
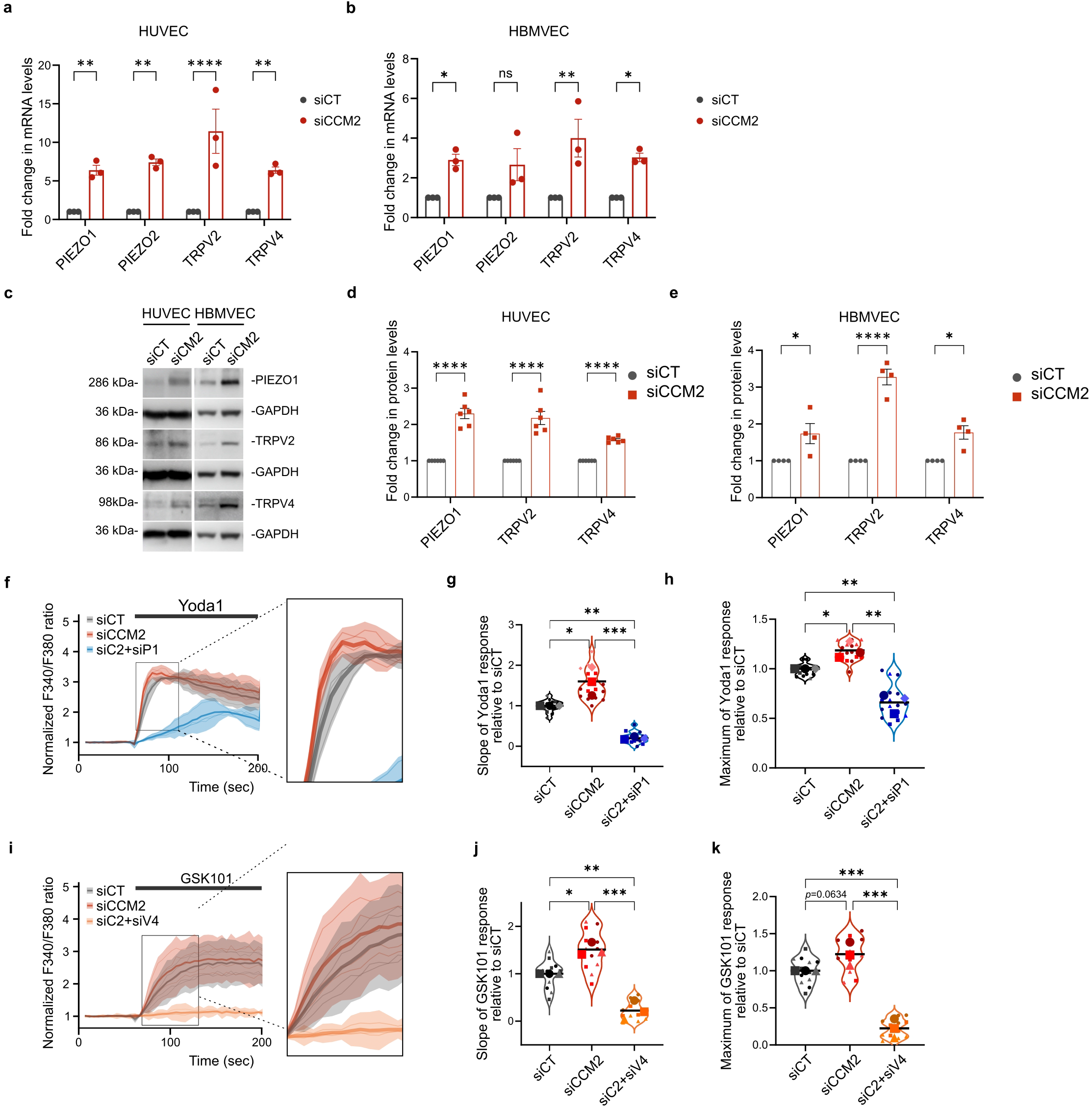
PIEZO1, TRPV2 and TRPV4 are overexpressed upon CCM2 loss in primary human EC from brain microvessels. **(a-b)** : PIEZO1, PIEZO2, TRPV2 and TRPV4 mRNA levels in primary HUVEC (a) and HBMVEC (b) treated with siRNA. (*n*=3 independent experiments). Dots are the mean of three technical replicates. Bars indicate mean ± SEM. Two-way ANOVA with Šídák’s correction. *p < 0.05, ****p < 0.0001. Exact p-values in (a) are PIEZO1: 0.0098 ; PIEZO2 : 0.0023; TRPV2: <0.0001; TRPV4: 0.0096. Exact p-values in (b) are PIEZO1: 0.0382; PIEZO2: 0.0811; TRPV2: 0.0011; TRPV4: 0.0254. **(c)** : Representative images of western blots of PIEZO1, TRPV2 and TRPV4 in CT- and CCM2-silenced HUVEC and HBMVEC. Pictures of GAPDH control for PIEZO1 and TRPV4 come from the same gel. **(d-e)** : PIEZO1, TRPV2 and TRPV4 normalized protein levels in HUVEC (d) and HBMVEC (e). For HUVEC : n=6 independent experiments. For HBMVEC: n=4 independent experiments. Bars indicate mean ± SEM. Multiple unpaired t-tests. *p < 0.05, ****p < 0.0001. Exact p-values in (d) are PIEZO1: 0,000003 ; TRPV2: 0,000063; TRPV4: <0,000001. Exact p-value in (e) are PIEZO1: 0.035738; TRPV2 : 0.00004 ; TRPV4 : 0.005529 **(f) (i):** Kinetics of calcium entry in HUVECs treated with siRNA, in response to PIEZO1 agonist Yoda1 (2.5 µM (f) or TRPV4 agonist GSK1016790A (25 nM). The graph shows normalized fluorescence intensity (F340/F380nm, I/I0) of Fura-2AM loaded HUVECs. Data are individual curve responses (4 technical replicates; thin lines) and mean (thick line) ± 95% confidence intervals (shading) from one experiment. The graph is representative of 3 independent experiments. **(g-h) and (j-k)** : Slope (g and j) and peak (h and k) of fluorescence curves from Fura-2AM loaded HUVECs treated with siRNA in response to Yoda1 or GSK1016790A, normalized to siCT. Small dots are technical replicates, bigger dots are mean for each biological replicates (n = 3 independent experiments). One-way ANOVA with Tukey’s correction. *p < 0.05, ****p < 0.0001. Exact p-values are: in g: 0.0257 (siCT-siCCM2), 0.0069 (siCT-siC2+siP1), 0.0004 (siCCM2-siC2+siP1); in h : 0.0469 (siCT-siCCM2), 0.0058 (siCT-siC2+siP1), 0.0011 (siCCM2-siC2+siP1); in j: 0.0104 (siCT-siCCM2), 0.0022 (siCT-siC2+siV4), 0.0003 (siCCM2-siC2+siV4); in k :0.0634 (siCT-siCCM2), 0.0007 (siCT-siC2+siV4), 0.0003 (siCCM2-siC2+siV4).

These data confirm that in vitro CCM models faithfully recapitulate the upregulation of PIEZO1, TRPV2, and TRPV4 observed in human CCM lesions, validating their use for mechanistic studies. Previous studies have demonstrated that the loss of the CCM complex compromises endothelial integrity and reactivates sprouting angiogenesis. To assess their common and specific roles in the different mechanical processes occurring during CCM progression, we selectively silenced each of these channels (Fig. S2) or inhibited their activity pharmacologically in CCM2-silenced background.

### PIEZO1 and TRPV4 disrupt CCM2-silenced endothelial integrity by weakening cell-cell junctions, while PIEZO1 and TRPV2 promote ECM adhesion and degradation

Loss of CCM2 compromises endothelial monolayer integrity by disrupting the balance between cell–cell and cell–ECM adhesions²⁻⁸. In the absence of the CCM1/2 complex, VE-cadherin-mediated junctions weaken, whereas integrin-dependent adhesions strengthen and acquire matrix-degradative properties, enabling these mutant ECs to invade the surrounding ECM ⁴⁻⁵,⁹. Given the roles of PIEZO1, TRPV2, and TRPV4 in adhesion, migration, and invasion in cancer cells, we investigated whether they are also involved in the adhesive and invasive defects of CCM2-mutant ECs.

We first assessed their role in cell–cell junction disruption. HUVEC monolayers silenced for siCCM2, and additionally silenced or not for each channel individually, were cultured for 48h on fibronectin-coated glass and stained for β−catenin (which clusters at VE-cadherin-based adherens junctions) and F-actin. CCM2 deficiency leads to thinner junctions (Fig. 3a)^12,14,26,27^, compared to the large overlapping junctions observed in control (siCT) cells. Automated segmentation and quantification showed that PIEZO1 silencing restored junction thickness to control levels (Fig. 3a, b), while TRPV4 silencing showed a trend toward restoration. TRPV2 and PIEZO2 had no restorative effect (Fig. 3a, b). These results are indicative of the primary role of overexpressed PIEZO1 in destabilizing cell-cell junction in siCCM2 ECs. We next examined the actin cytoskeleton. In control ECs, a cortical actin rim supports VE-cadherin junctions (Fig. 3a). In CCM2-silenced ECs, this rim is reduced in favor of cytosolic transversal stress fibers^14^ (Fig. 3a). Additional silencing of TRPV4 significantly restored the cortical actin rim as measured by the ratio between cortical versus cytosolic F-actin (Fig. 3a, c). Surprisingly, this was not the case when PIEZO1 was silenced, with transversal stress fibers still being observed (Fig. 3a, c). This revealed an unusual decoupling between the perturbation of VE-cadherin dependent cell-cell junctions and actin cytoskeleton remodeling.

**Figure 3:**
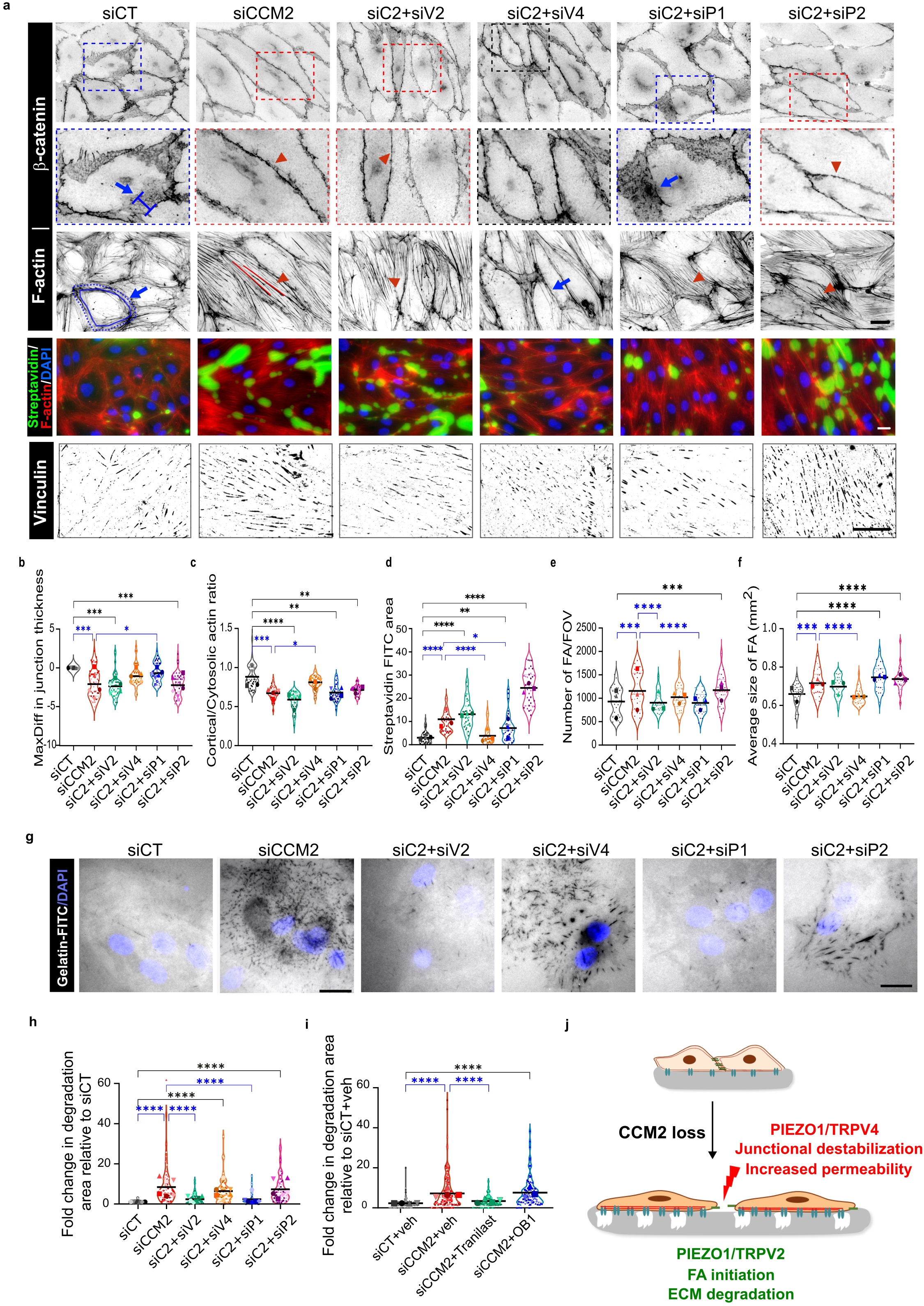
Overexpressed TRPV2 and TRPV4 combine with PIEZO1 to destabilize CCM2-silenced endothelial monolayers. **(a)** : Representative images of silenced HUVECs monolayers stained for cell-cell junctions (β-catenin, upper panel), F-actin (phalloidin, third panel), junctional permeability (merged streptavidin-FITC with F-actin and DAPI) and cell-ECM adhesions (vinculin, lower panel). Second panel shows magnified views of boxed regions of β-catenin images. Blue arrows and brackets delineate thick cell-cell junctions and red arrowheads point at thin disrupted junctions. In the F-actin panel, blue arrow and circles point at cortical organization of F-actin whereas red lines and arrowheads point at cytosolic transversal actin stress fibers. In the Streptavidin/F-actin/DAPI panel, green fluorescence depicts areas permeable for FITC-labeled streptavidin. Abbreviations: C2:CCM2; V2:TRPV2; V4:TRPV4. P1:PIEZO1; P2: PIEZO2. Scale bar 20µm. **(b-i) :** Small dots are technical replicates per condition, bigger dots are mean for each biological replicate. **(b)** : Maximal difference (MaxDiff) in cell-cell junction thickness. Negative values correspond to thinner junctions than CT HUVECs. Technical replicates (10-24 fields of view (FOV) per condition, approximately 70 cells per FOV) from 5 independent experiments. Two-way ANOVA with Šídák’s correction for multiple comparisons. *p < 0.05, ****p < 0.0001. Exact p-values are 0.0008 (siCT vs siCCM2), 0.0002 (siCT vs siC2+siV2), 0.0004 (siCT vs siC2+siP2), 0.0321 (siCCM2 vs siC2+siP1). **(c)** : Actin intensity ratio between cortical and cytosolic cell area. Technical replicates (8-14 FOV per condition, each contains approximately 70 cells) from 4 independent experiments. Two-way ANOVA with Šídák’s correction for multiple comparisons. *p < 0.05, ****p < 0.0001. Exact p-values are 0.0009 (siCT-siCCM2), <0.0001 (siCT-siC2+siV2 and siCT-siC2+siP1), 0.0002 (siCT-siC2+siP2), 0.0285 (siCCM2-siC2+siV4). **(d)** : Measurements of endothelial monolayer permeability expressed as the percentage of FITC-stained area per FOV. Technical replicates (10-14 FOV per condition, each contains approximately 10-20 cells) from 3 independent experiments. Two-way ANOVA with Šídák’s correction for multiple comparisons. *p < 0.05, ****p < 0.0001. Exact p-values are <0,0001 (siCT-siCCM2; siCT-siC2+siV2; siCT-siC2+siP2; siCCM2-siC2+siV4), 0.0014 (siCT-siC2+siP1), 0.0248 (siCCM2-siC2+siP1). **(e)** : Number of vinculin-stained focal adhesions. Technical replicates (10 FOV per condition, each contains between 5-10 cells) from 3 independent experiments. Two-way ANOVA with Tukey’s correction for multiple comparisons. *p < 0.05, ****p < 0.0001. Exact p-values are 0.0006 (siCT-siCCM2), 0.0069 (siCT-siC2+siP2), <0.0001 (siCCM2-siC2+siV2 and siCCM2-siC2+siP1). **(f)** : Average size of focal adhesions. Technical replicates (10 FOV per condition, each contains between 5-10 cells) from 3 independent experiments. Two-way ANOVA with Tukey’s correction for multiple comparisons. *p < 0.05, ****p < 0.0001. Exact p-values are 0.001 (siCT-siCCM2), < 0.0001 (siCT-siC2+siP1 and siCT-siC2+siP2), <0.0001 (siCCM2-siC2+siV4). **(g)** : Representative images of fluorescent gelatin after overnight culture of silenced HUVECs (nuclei in blue). Dark stripes are matrix degradation areas. Scale 20µm. **(h)** : Fold change in matrix degradation area relative to siCT. Technical replicates (at least 6 FOV per condition) from 6 independent experiments. Two-way ANOVA with Tukey’s correction for multiple comparisons. *p < 0.05, ****p < 0.0001. Exact p-values are < 0.0001 (siCT vs siCCM2), < 0.0001 (siCT vs siC2+siV4 and siCT vs siC2+siP2), < 0.0001 (siCCM2 vs siC2+siV2 & siCCM2 vs siC2+siP1). **(i)** : Quantification of matrix degradation area by siCCM2 HUVEC treated with Tranilast (TRPV2 inhibitor) or OB1 (PIEZO1 inhibitor). Technical replicates (at least 25 FOV per condition) from 4 independent experiments, except for siCCM2+OB1 (n=3). Two-way ANOVA with Tukey’s correction for multiple comparisons. *p < 0.05, ****p < 0.0001. Exact p-values are < 0.0001 (siCT+veh vs siCCM2+veh and siCT+veh vs siCCM2+OB1), < 0.0001 (siCCM2+veh vs siCCM2+Tranilast). **(j)** : Proposed role of mechanosensitive channels in the destabilization of the CCM2-mutant endothelial monolayer.

Since morphologically abnormal cell-cell junctions are functionally linked with increased endothelial permeability^14,26,27^, we evaluated the role of each MSC in regulating the passage of fluorescent streptavidin (52 kDa) across the monolayer as a proxy for permeability measurements^28^. Consistently with their restorative effect on the morphology of VE-cadherin junctions, PIEZO1 and TRPV4 silencing significantly reduced the passage of FITC-streptavidin across the cell-cell junctions while TRPV2 and PIEZO2 silencing did not (Fig. 3a, d). TRPV4 fully restored the endothelial barrier while PIEZO1 restored it only partially (Fig3 d) indicating that the tightness of cell-cell junctions relies more strongly on their underlying cortical actin cytoskeleton than on VE-cadherin-dependent membrane borders.

Thus, PIEZO1 and TRPV4 cooperate to provoke junctional permeability defects in CCM2-mutant ECs, with PIEZO1 acting on VE-cadherin junctions and TRPV4 regulating their underlying actin cytoskeleton.

We then evaluated the contribution of these channels to increased cell–ECM adhesion which was previously found to counterbalance the weakening of cell-cell junctions upon CCM loss^12^. CCM2 silencing led to more and larger vinculin-stained ECM adhesions across the EC monolayer (Fig. 3. a, e, f), a phenotype we previously observed in migratory sparse CCM1/2-silenced ECs^12^. PIEZO1 and TRPV2 silencing both reduced the number of adhesions to control levels (Fig. 3e), while TRPV4 silencing restored their size (Fig. 3f). PIEZO2 silencing had no restorative effect. These results suggest that PIEZO1 and TRPV2 may be required for focal adhesion initiation, whereas TRPV4 promotes their maturation, likely through the remodeling of the actin cytoskeleton into stress fibers (Fig. 3a, c).

Silencing CCM2 in ECs not only enhances their adhesion to the ECM but also promotes its degradation, thereby sustaining their invasive phenotype ^13,17^. When ECs are cultured overnight on fluorescent gelatin, focal areas of degradation can be observed, which shapes resembled cell-ECM adhesion footprint. In line with our previous observations^13^, siCCM2 ECs degraded the gelatin, whereas CT ECs barely did (Fig. 3g). TRPV2 and PIEZO1 additional silencing both completely blocked degradation (Fig. 3g, h) consistent with their capacity to restore the number of cell-ECM adhesions to CT level (Fig. 3e). In contrast, TRPV4 or PIEZO2 silencing had no restorative effect on ECM degradation. We previously showed that siCCM2 HUVECs overexpress numerous ECM remodelers, including MMPs that are responsible for ECM degradation^13^, among which MMP14 (MT1-MMP) that activates MMP9 at cell-ECM adhesive sites. Interestingly, increased MMP14 and MMP9 expression in CCM2-silenced ECs was restored to CT expression levels upon TRPV2 silencing and to a lesser extent upon PIEZO1 silencing (Fig. S3a, b), indicating an additional transcriptional role of these channels. To evaluate the effect of pharmacological inhibition of these MSCs on ECM degradation, we tested the effect of Tranilast, a TRPV2 inhibitor, and of OB-1, an indirect inhibitor of PIEZO1^29^. Consistent with the restoration upon TRPV2 silencing, Tranilast completely inhibited gelatin degradation by CCM2 depleted cells, whereas OB-1 did not (Fig. 3i).

These findings reveal complementary roles for TRPV4 and TRPV2 in CCM2-mutant ECs: TRPV4 destabilizes cell–cell junctions and increases barrier permeability, while TRPV2 enhances the formation of ECM adhesions and ECM degradation. Moreover, these findings unravel complementary roles between PIEZO1 and TRPV4 at intercellular junctions, and overlapping roles between PIEZO1 and TRPV2 at cell–ECM adhesion sites (Fig. 3j). While PIEZO1–TRPV4 interaction has been previously documented, with PIEZO1 activating TRPV4, our data uncover a novel functional coupling between PIEZO1 and TRPV2, not previously reported.

### TRPV2 stimulates 3D-invasion by CCM2-silenced angiogenic tip cells by favouring filopodia formation

In addition to their impaired ability to form stable intercellular junctions and their enhanced cell-ECM adhesion, CCM2 mutant ECs acquire a hyper-angiogenic tip cell phenotype. Using a 3D degradable polyethylene glycol (PEG) - based angiogenic sprouting assay compatible with 3D traction force microscopy (TFM)^30^ (Fig. 4a) we previously showed that CCM2 mutant HUVECs actively invade 3D-PEG gels (Fig. 4a)^13,17^. They form numerous filopodia at their leading tip to exert pulling forces on the matrix, while degrading it at the tip cell rear to facilitate motion of the cell body^17^. Consistent with these results, siCCM2 HBMVECs invaded more deeply into the gel compared to siCT HBMVECs (Fig. 4a, c). Additional silencing of TRPV2 significantly blocked invasion by siCCM2 ECs, reverting the invasion distance to that of siCT level (Fig. 4a, c), an effect confirmed upon inhibition of TRPV2 with Tranilast (Fig. 4d). This reduced invasion correlated with a reduced number of tip cell filopodia (Fig. 4b, e) and lowered F-actin intensity to that of siCT sprouts (Fig. 4f). These findings agree well with the previously uncovered role of TRPV2 in enhancing neurite outgrowth^31,32^. Additional silencing of PIEZO1 showed a trend towards restoration of invasion distance and filopodia number (Fig. 4c, e) and a significant decrease of the F-actin intensity with respect to siCCM2 HBMVECs (Fig. 4f). Strikingly, additional TRPV4 silencing increased invasion even further compared to siCCM2 HBMVECs (Fig. 4a, c).

**Figure 4:**
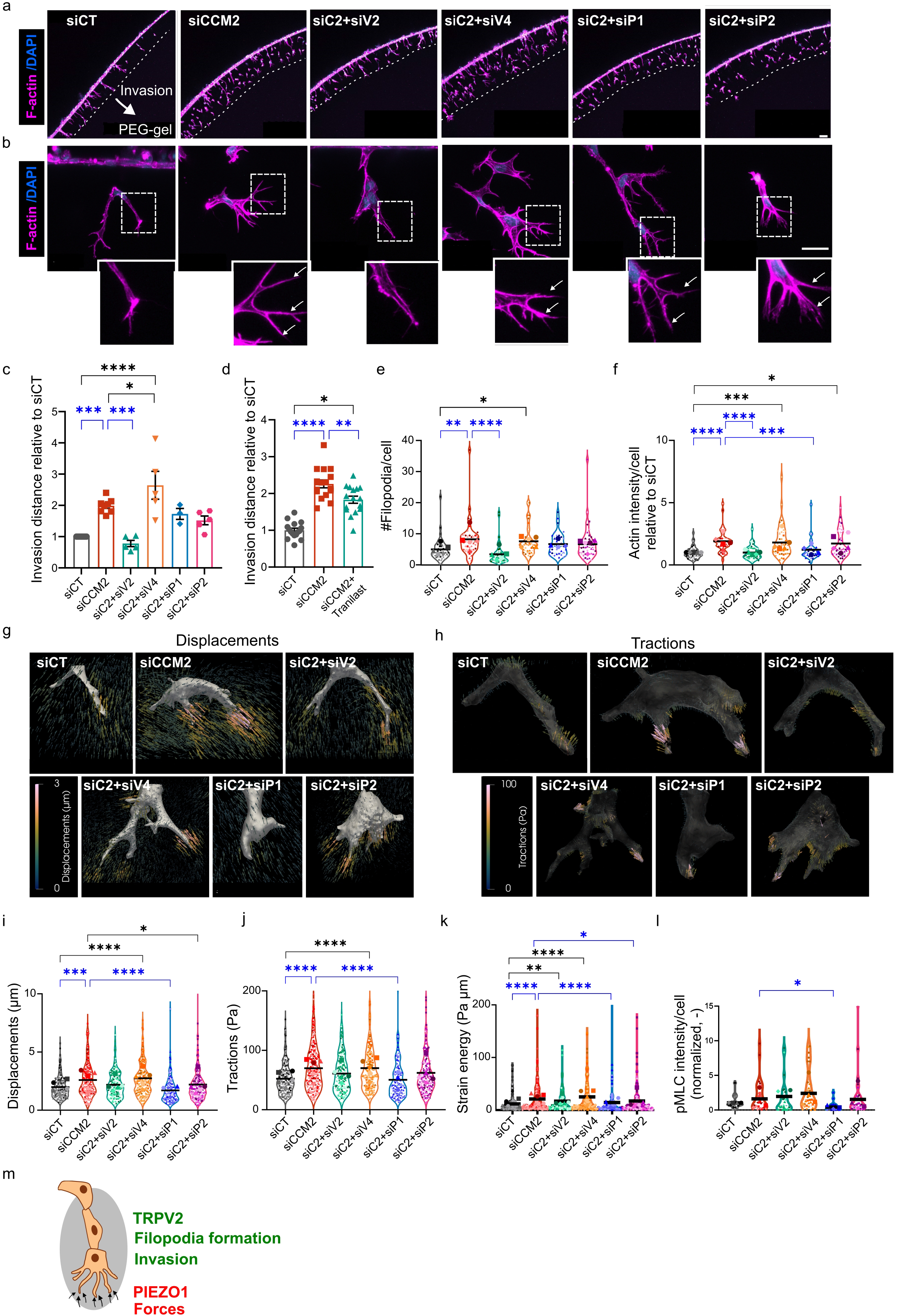
PIEZO1 and TRPV2 promote the increased invasiveness of CCM2-silenced ECs. **(a)** and **(b):** Maximum F-actin intensity projections of the entire monolayer of transfected HBMVECs **(a)** and leading tip regions **(b)** after 18h of invasion. Dash line indicates the invasion front. White arrows mark filopodia. Scale bars indicate 200µm and 30µm respectively. **(c) :** Maximum invasion distance. siCT & siCCM2 n=8, siC2+siP1 n=3, siC2+siP2 & siC2+siT2 & siC2+siT4 from 5 independent experiments. Bars indicate mean ± SEM. One-way ANOVA with Tukey’s multiple comparisons test. Exact p-values are 0.0002 (siCT vs siCCM2), <0.0001 (siCT vs siC2+siV4), 0.0001 (siCCM2 vs siC2+siV2), 0.0485 (siCCM2 vs siC2+siV4). **(d) :** Maximum invasion distance upon treatment with Tranilast. Technical replicates (8 FOVs) from 2 independent experiments. Bars indicate mean ± SEM. One-way ANOVA with Tukey’s multiple comparisons test. Exact p-values are <0.0001 (siCT vs siCCM2), 0.0218 (siCT vs siCCM2+Tranilast), p=0.0012 (siCCM2 vs siCCM2+Tranilast). **(e) :** Number of filopodia normalized to that of nuclei. Technical replicates (5 to 10 sprouts) from 5 independent experiments except for siC2+siP1 (n = 4). One-way ANOVA with Kruskal-Wallis test. Exact p-values are 0.0031 (siCT vs siCCM2), 0.0411 (siCT vs siC2+siV4), <0.0001 (siCCM2 vs siC2+siV2). **(f) :** Total F-actin intensity normalized to that of nuclei. Technical replicates (5 to 10 sprouts) from 5 independent experiments except for siC2+siP1 (n = 4). One-way ANOVA with Kruskal-Wallis test. Exact p-values are <0.0001 (siCT vs siCCM2), 0.0003 (siCT vs siC2+siV4), 0.0142 (siCT vs siC2+siP2), <0.0001 (siCCM2 vs siC2+siV2), 0.0005 (siCCM2 vs siC2+siP1). **(g-h)** : Representative 3D renders of displacements and tractions around angiogenic sprout tips. **(i-k) :** Quantification of 99th percentile bead displacements , tractions and strain energy around sprout tips. Technical replicates (19 to 70 sprout tips) from 5 independent experiments except for siC2+siP1 (n = 3). One-way ANOVA with Kruskal-Wallis test. For the displacements (**i**), exact p-values are 0.0003 (siCT vs siCCM2), <0.0001 (siCT vs siC2+siV4 & siCCM2 vs siC2+siP1), 0.0155 (siCCM2 vs siC2+siP2). For the tractions (**j**), exact p-values are <0.0001 (siCT vs siCCM2 & siCT vs siC2+siV4 & siCCM2 vs siC2+siP1). For the strain energy (**k**), exact p-values are <0.0001 (siCT vs siCCM2 & siCT vs siC2+siV4 & siCCM2 vs siC2+siP1), 0.0037 (siCT vs siC2+siV2), 0.0348 (siCCM2 vs siC2+siP2). **(l) :** pMLC intensity normalized to nuclei number. Technical replicates (5 to 10 sprouts) from 4 independent experiments except for siC2+siP1 (n = 3). One-way ANOVA with Kruskal-Wallis test. The exact p-value is 0.0128 (siCCM2 vs siC2+siP1). **(m) :** Proposed roles of mechanosensitive channels in sprouting invasion by CCM2-mutant ECs.

Using 3D TFM, we previously reported that branchy CCM2-silenced HUVECs exert stronger, ROCK-dependent pulling forces on their surrounding matrix during invasion^17^. This was also observed for siCCM2 HBMVECs, which generated larger matrix displacements, tractions and strain energy compared to siCT HBMVECs (Fig. 4g-k). PIEZO1 was the only channel that upon additional silencing led to a significant reduction of all three metrics of cell-ECM mechanical interactions (Fig. 4g-k). Analysis of phosphorylated Myosin Light Chain (pMLC) staining further confirmed PIEZO1 as a key player in cell contractility, as its additional silencing resulted in restoring pMLC intensity to siCT levels (Fig. 4l).

We previously demonstrated that the hyper-angiogenic behaviour of siCCM2 ECs could be associated to the combined upregulation of F-actin rich filopodia, actomyosin contractility and ECM degradation^13,17^. When combining our 2D gelatin degradation and 3D invasion data, we can witness overlapping as well as complementary roles of TRPV2 and PIEZO1 overexpression in controlling features that contribute to siCCM2 ECs hyper-angiogenic tip cell phenotype (Fig. 4m): (i) degradation and actin intensity are under the control of both TRPV2 and PIEZO1, (ii) filopodia number is controlled by TRPV2 only and (iii) cell-ECM tractions and pMLC intensity are controlled by PIEZO1 only. Interestingly, only TRPV2 silencing reduced invasion distance to siCT levels, which suggests that from all investigated features, invasive behaviour is best predicted from filopodia number.

Integration of these *in vitro* data with our patient analyses provides a coherent molecular explanation for the predictive correlation between the TRPV2:TRPV4 ratio and the size of CCM lesions. Indeed, we uncover a functional coupling of PIEZO1 with TRPV4 at cell–cell junctions and with TRPV2 at cell–ECM adhesion sites. Mechanistically, PIEZO1 and TRPV4 destabilize intercellular junctions in a complementary way, by affecting VE-cadherin and actin organization respectively, while PIEZO1 and TRPV2 jointly promote ECM adhesion, degradation, and invasive behavior. Consequently, an elevated TRPV2:TRPV4 ratio could be associated with TRPV2-dependent endothelial invasion into the brain parenchyma, as a way to increase lesion size, at sites of TRPV4-dependent barrier disruption. Notably, this ratio emerges as a potential composite biomarker for CCM lesion progression.

### Reduction of Trpv4 signalling alleviates cardiovascular defects in ccm2 mutants

Having characterized the molecular and cellular processes driven by overexpressed TRPV channels by means of *in vitro* monolayer and angiogenic invasion models, we sought to further evaluate their functional relevance in an *in vivo* CCM model. To this end, we used ccm2 mutant zebrafish embryonic hearts to analyze hallmark features of the CCM phenotype. In zebrafish, there is no expression of a Trpv2 channel but expression of an ortholog which has functional similarities to TRPV1 but not TRPV2^33^. However, an ortholog of TRPV4 exists. So, we decided to lower its activity in ccm2 mutant zebrafish embryos through genetic depletion and pharmacological inhibition. In this ccm model, cardiac ballooning arises from the loss of the endocardial atrioventricular canal (AVC)^34^ (Fig. 5 ai-ii).

**Figure 5:**
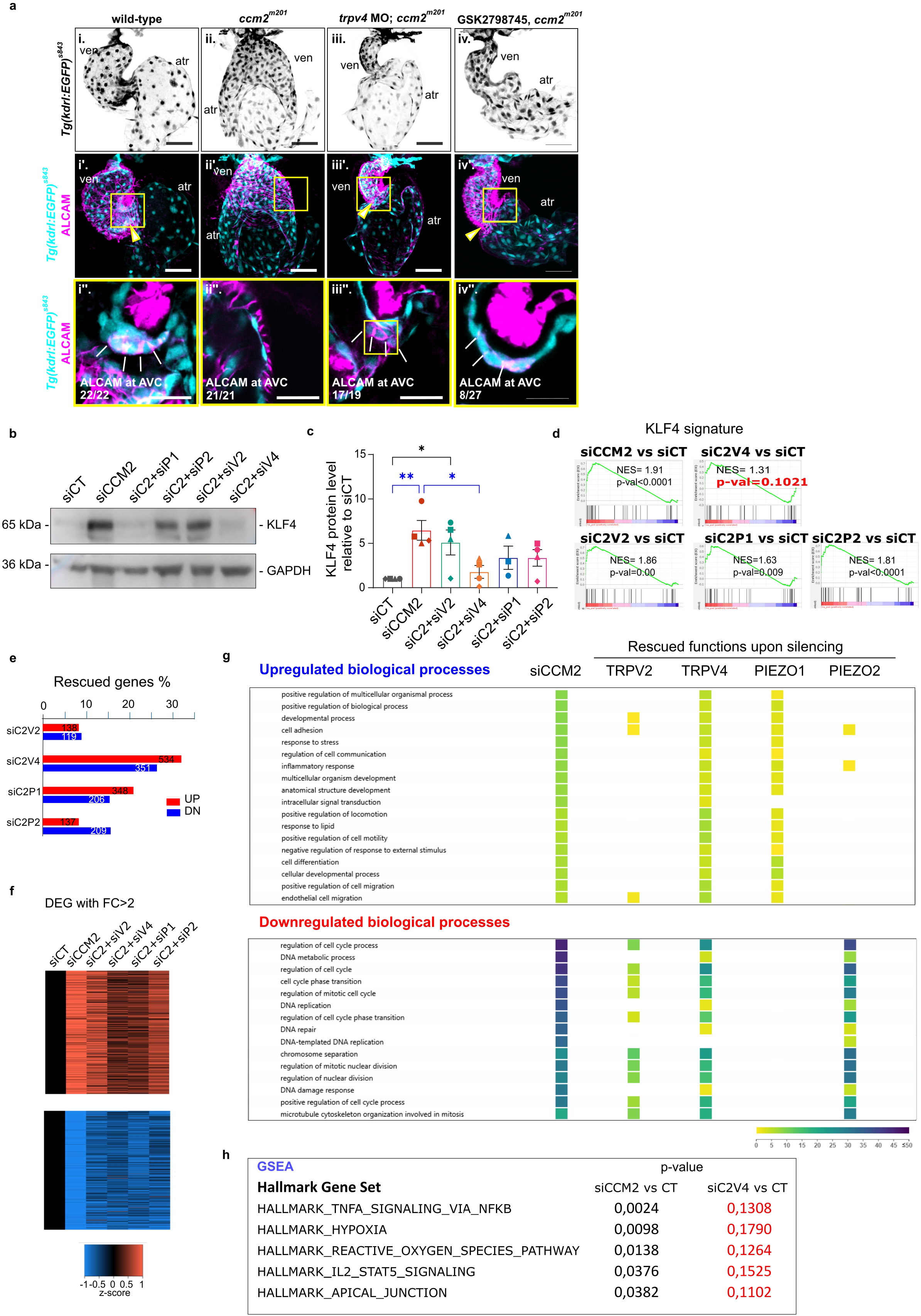
TRPV4 inhibition rescues ccm2 zebrafish cardiac defects, KLF4 upregulation and transcriptomic dysregulation. **(a) :** Endocardium at 56 hpf, marked *by Tg(kdrl:EGFP)^s843^* expression in wild-type (i), *ccm2^m201^* (ii), and trpv4 MO-injected *ccm2^m201^* (iii), and GSK2798745-treated *ccm2^m201^* (iv). The zebrafish endocardium forms the atrioventricular canal (AVC, arrowhead) between the ventricle and the atrium at 56 hpf (i’). Endocardial cells of the AVC are marked by the expression of ALCAM at cell-cell junctions (i’’; white lines). In *ccm2^m201^* mutants, there is no AVC constriction (ii’) and endocardial cells lack ALCAM expression (ii’’). The endocardial defects in zebrafish *ccm2^m201^* mutants are suppressed when Trpv4 is genetically depleted (*trpv4* MO; *ccm2^m201^*) (iii). *trpv4* MO; *ccm2^m201^* mutants form an AVC including restored cuboidal endocardial cell shape and ALCAM expression at cell-cell junctions (iii’,iii’’). Similarly, *ccm2^m201^* mutants treated with trpv4-inhibitor GSK2798745 (iv) show a restoration of the AVC constriction and ALCAM expression at cell-cell junction (iv’, iv’’). Number in the lower panel correspond to the ratio of embryos with WT phenotype. **(b) :** Representative images of western blot of KLF4 in silenced HBMVEC. GAPDH is shown as a loading control. **(c)** : KLF4 normalized protein levels detected by western blot in HBMVEC. n=3 independent experiments. Bars indicate mean ± SEM. One-way ANOVA with Tukey’s correction. *p < 0.05, ****p < 0.0001. Exact p-values are 0.0062 (siCT vs siCCM2); 0,0104 (siCT vs siC2+siV2); 0.036 (siCT vs siC2+siV4). **(d)** : Gene set enrichment analysis of KLF4 signature (from ^73^) in the transcriptome of silenced HUVECs using GSEA software from Broad Institute UC San Diego. NES: Normalized enrichment score. **(e)** : Percentage of dysregulated expressed genes (DEG) (p-value <0.05) in CCM2-silenced transcriptome rescued upon additional silencing of the MSCs. **(f)** : Comparison heatmap of the expression level of CCM2-dependent DEG with fold change (FC)>2, p<0.05 among the different silencing conditions. **(g)** : Gene ontology analyses using G-profiler of enriched biological processes associated with DEG (FC>2, p<0.05) in siCCM2 HUVECs and their rescue by additional silencing of the MCSs. **(h)** : Gene set enrichment of human curated gene sets (C2 collection, GSEA Broad Institute UC San Diego) in the transcriptome of silenced HUVECs. Hallmark gene sets uniquely rescued by TRPV4 additional silencing are shown.

We injected an antisense morpholino oligo targeting trpv4 into embryos from *ccm2^m201^; Tg(kdrl:EGFP)^s843^* heterozygous parents. In these embryos, endocardial cells express GFP and can therefore be followed over time. At 54-56 hours post-fertilization (hpf), Trpv4 knockdown suppressed *ccm2^m201^* mutant cardiac defects by rescuing the formation of the AVC (Fig. 5a-iii.). The morphogenetic changes normally occurring in endocardial cells during AVC formation were restored, such as acquired cuboidal shape (Fig. 5a-iii”) and regained cell-cell junctions as evidenced by junctional Activated Leukocyte Cell Adhesion Molecule (ALCAM) staining (Fig. 5a-iii”). Similarly, treatment of embryos from the 16-somite stage to 54–56 hpf with GSK2798745, a selective TRPV4 channel inhibitor, also suppressed *ccm2^m201^* mutant cardiac defects (Fig. 5a-iv-iv”). This included a restoration of the AVC with ALCAM expression at intercellular junctions and changes in cell shape (Fig. 5iv”). For comparison, the effects of Trpv4 knock-down or GSK2798745 on wild-type embryos are shown in Fig. S4. They did not affect the morphology of the developed heart.

With its inhibition leading *in vivo* to regained cell-cell junctions as observed *in vitro*, these results confirm the role of TRPV4 in disrupting the endothelial barrier integrity and reinforce considering the use of its pharmacological inhibitors in clinics.

### TRPV4 upregulates KLF4 expression and drives CCM2-silenced EC reprogramming

Previous studies using zebrafish CCM models have been instrumental in uncovering the causative role of the mechanosensitive transcription factors KLF2 and KLF4 in the cell state transitions of CCM-deficient ECs. Notably, they showed that their downregulation can rescue heart development by restoring AVC formation^35,36^. These findings were further supported by studies in mouse CCM models, which confirmed a role for these transcription factors in CCM lesion formation in mammals^37,38^.

We therefore hypothesized that overexpressed TRPV4 might not only disrupt cell-cell junctions but also contribute to the dysregulation of the transcriptional program triggered by KLF2 and 4 overexpression. Indeed, silencing TRPV4 (and, to a lesser extent, PIEZO1) prevented KLF4 protein overexpression in CCM2-silenced HBMVECs (Fig. 5b, c) and HUVECs (Fig. S5a, b) demonstrating TRPV4 central role in controlling KLF4 expression. To investigate further its contribution to the transcriptomic reprogramming of CCM2-silenced ECs, we performed bulk RNA sequencing of siCCM2 HUVECs, either alone or in combination with the silencing of each channel. In line with the reduction in KLF4 protein, the enriched KLF4 signature in siCCM2 HUVECs was selectively abolished upon additional TRPV4 silencing (Fig. 5d), confirming that TRPV4 overexpression specifically drives KLF4 transcriptional hyperactivation in CCM2-mutant ECs. Similarly, the enrichment in the KLF2 signature was also specifically lost following additional TRPV4 silencing (Fig. S5c). More broadly, we observed that the additional silencing of TRPV4 normalized the expression of around 30% of the up- and down-regulated genes (DEGs) upon CCM2 loss, compared to only 10–20% for the other channels (Fig. 5e, f) (Table S2). This normalization was potent enough to rescue most of the dysregulated biological processes associated with the loss of CCM2 (Fig. 5g). This includes upregulated cell adhesion, migration or inflammatory response. This effect was only replicated by silencing PIEZO1, which however had no effect on downregulated biological processes. Additionally, TRPV4 silencing rescued a broad set of downregulated processes which we previously characterized^13^, such as cell cycle, DNA replication, and chromosome segregation, a capacity shared with PIEZO2 silencing which however had no effect on upregulated biological processes. Gene set enrichment analysis confirmed that only TRPV4 silencing abolished enrichment in CCM hallmarks such as hypoxia, ROS pathways, TNFα and IL2 signaling, and apical junction dysfunction (Fig. 5h).

Collectively, these findings establish TRPV4 as a central regulator of transcriptomic reprogramming in CCM2-mutant ECs, consistent with evidence that junctional signaling drives CCM gene dysregulation^39,40^.

## Discussion

While most research has focused on the intracellular signaling pathways triggered by mutations in CCM1-3, PIK3CA, or MAP3K3 genes that drive lesion formation^41^, the contribution of environmental cues has remained largely overlooked. Yet, external mechanical signals combine with EC genetic defects for the formation of CCM lesion in low-flow blood vessels^4,5,18^. Here, we reveal that the ECs lining human CCM lesions overexpress mechanosensors responsive to fluid shear stress and ECM stiffness. Following an unprecedented integrative approach, we further demonstrate that the identified PIEZO1, TRPV2 and TRPV4 mechanosensors act in concert to drive characteristic CCM endothelial defects in cellular and animal models. We reveal a functional coupling between PIEZO1 and TRPV4 at intercellular junctions, and between PIEZO1 and TRPV2 at cell–ECM adhesion sites. Mechanistically, PIEZO1 and TRPV4 destabilize intercellular junctions in a complementary way, by affecting VE-cadherin and actin organization respectively, while PIEZO1 and TRPV2 promote ECM adhesion, degradation, and invasive behavior (Fig. 6). Strikingly, the correlation between TRPV2:TRPV4 expression ratio and the size of the lesions is supported by our mechanistic data. Indeed, elevated TRPV2:TRPV4 ratio would result in TRPV2-dependent endothelial invasion into the brain parenchyma at sites of TRPV4-dependent barrier disruption, favoring CCM lesion expansion at sites of CCM onset.

**Figure 6:**
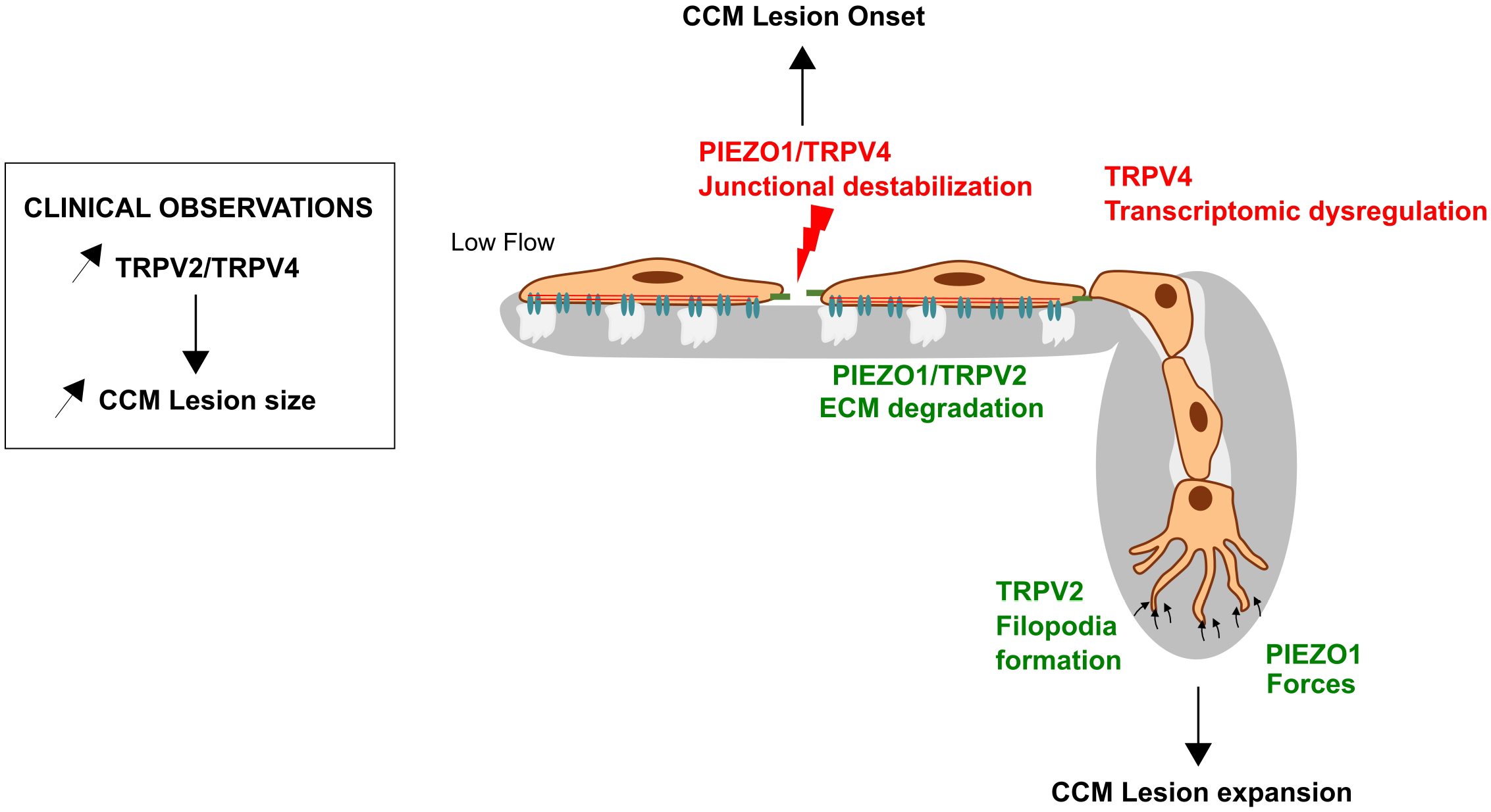
Proposed model for the complementary roles of PIEZO1, TRP V2 and TRPV4 in CCM onset and expansion. PIEZO 1 and TRPV4 complementarily destabilize intercellular junctions by affecting VE-cadherin and actin organization respectively, while PIEZO 1 and TRPV2 promote ECM adhesion, degradation, and sprouting angiogenesis. Based on these mechanistic findings, the correlation between the TRPV2:TRPV4 expression ratio and the brainstem lesion volume may result from the TRPV2-dependent endothelial invasion into the brain parenchyma at sites of TRPV4-dependent barrier disruption favoring CCM lesion expansion at sites of CCM onset.

Findings advocate for an early and broad role of biomechanical stimulation over lesion lifespan. *In vivo* zebrafish models have shown that blood flow mitigates the appearance of vascular defects^4,18,42^. In patients, low perfusion within existing lesions can predict bleeding in the year following imaging^6^. Mechanistically, CCM-silenced ECs are not able to adapt to low shear stress *in vitro* and behave as if they were exposed to high shear stress^5^. Although the contributions of mechanosensitive effectors, including ROCK1/2 and KLF2/4, to CCM pathology are well documented, the upstream mechanosensors regulating the integrin- and VE-cadherin-dependent dysregulated cellular responses remained elusive. Our identification of mechanosensitive channels as key players is particularly compelling, considering their essential roles in vascular stability and angiogenesis.

In the brain, PIEZO1 and TRPV4 control the blood brain barrier in response to mechanical signals^43–45^. PIEZOs are pivotal force sensors of endothelial flow with a pleitropic role in vascular development and homeostasis^46^. PIEZO1 is a primary force sensor directly activated by the stretching of the plasma membrane upon fluid shear stress. Its activation triggers TRPV4 downstream opening through activation of Phopholipase A2^47^. PIEZO1 interacts with the mechanosensory complex containing VE-cadherin and PECAM to tailor junctional structure to mechanical requirements^48,49^. Excessive activation of PIEZO1 results in calpain-induced disruption of VE-cadherin adhesions^50–52^. Consistently, we found that the thinning of VE-cadherin borders in siCCM2 ECs is dependent on overexpressed PIEZO1 which does not affect the underlying actin cytoskeleton, an effect dependent on overexpressed TRPV4. The actin remodeling activity of TRPV4 has been well documented and relies on the control of the RhoA/ROCK pathway^53,54^. Indeed, activation of TRPV4 releases its inhibitory interaction with RhoA, which is then in turn activated by TRPV4-dependent Ca2+ influx^54^.

Going further, we demonstrate that in CCM mutant ECs, TRPV4 is responsible for the overexpression of KLF2 and KLF4. Its silencing restored AVC formation during ccm2 mutant cardiac development. This aligns with the reported modulation of klf2a expression by TRPV4-mediated Ca^2+^ waves during zebrafish heart valve formation^55^. It is likely that a sustained TRPV4 activation in the developing ccm2 mutant heart leads to dysregulated expression of KLF2 and subsequent defects in AVC formation. We demonstrated that TRPV4 has a primary role in the transcriptomic defects of CCM2-silenced ECs. It is striking to note that TRPV4 is overexpressed in brain ECs from Ccm3 KO mice (https://edgroup.shinyapps.io/MapCcm3EC/). Its expression is restricted to the venous capillary cluster at the origin of CCM3 lesions^23^ providing a crucial additional cue of its importance in CCM lesion onset.

Among the three MSCs that we identified, PIEZO1 only has so far been associated with vascular malformations while hereditary TRPV4 mutations are associated with neuromuscular and skeletal disorders^56^. A brief report identified mutations on PIEZO1 gene in a family with hereditary CCM^57^. PIEZO1 was shown to be overexpressed in Hereditary Hemorrhagic Telangiectasia (HHT) where it is involved in the signaling defects downstream of Alk1 loss-of-function^58^. HHT are arteriovenous malformations that also depend on the level of fluid shear stress but in a reversed manner compared to CCM. Mechanistically, in HHT, PIEZO1 overexpression overactivated PI3K and ERK5-KLF4 signaling pathways^58^. Besides KLF4 signaling, TRPV4 has also been shown to regulate PI3K/AKT signaling^59^. Therefore, it would be interesting to investigate whether the overactivation of PI3K and ERK in CCM results from PIEZO1 and TRPV4 overexpression. This could provide an explanation for cases where no PI3K gain-of-function mutations have been identified in CCM patients.

High-resolution 3D reconstruction of two human CCM brainstem lesions revealed a centrifugal pattern of microvascular buds and cavernous spaces originating from the feeding vessel and infiltrating the adjacent parenchyma^60^. The reactivation of an angiogenesis program in the CCM mutant ECs^61^ results in an excessive invasion of the surrounding tissue, a process that can be reproduced *in vitro*^30^. We previously showed that CCM2-silenced HUVECs have acquired a highly invasive ROCK-dependent tip cell phenotype which relies on filopodia formation, force exertion and ECM degradation^17^. Numerous and branchy filopodia generate strong tractions at the tip cell front. Here, we demonstrated that PIEZO1 and TRPV2 cooperatively trigger these events. They both stimulate ECM degradation, the first step of sprouting angiogenesis, most likely by increasing cell-ECM adhesion and MMP expression. In melanoma, TRPV2 activity is sufficient to confer invasiveness to melanoma cells by localizing in dynamic nascent adhesions at their leading edge, where it regulates Ca^2+^-mediated activation of calpain and the ensuing cleavage of talin, along with cofilin-dependent F-actin organization^62^. Consistently, silencing TRPV2 lowered the number of focal adhesions in CCM2-silenced EC monolayers suggesting a role in focal adhesion initiation. During invasion, silencing TRPV2 restores the number of filopodia in CCM2-silenced tip cells. This is consistent with its stimulating activity on neurite formation, extension and branching in neurons^31,32^. While PIEZO1 is well established as a direct mechanosensor, whether it generates forces remained poorly explored. Here we showed that PIEZO1 controls force exertion by the TRPV2-dependent filopodia in CCM2-silenced tip cells, an activity correlated with increased pMLC intensity.

These *in vitro* mechanistic findings provide a coherent rationale for our molecular and clinical observations in patients linking MSCs expression to lesion size and location. Brainstem lesions exhibited significantly higher PIEZO1 levels compared to those in other brain regions. Given that the brainstem is a confined and mechanically challenged region with the highest stiffness compared to the other regions of the brain^25^, these biomechanical constraints may drive PIEZO1 upregulation as part of an adaptive response. TRPV2 expression positively correlated with lesion size, a correlation that was found to be even more robust in the brainstem when considering the TRPV2:TRPV4 ratio. In melanoma, TRPV2 overexpression correlates with advanced malignancy and aggressive metastatic behavior^62^. Consistently, TRPV2 appears to drive cancer cell invasion and metastasis rather than proliferation across multiple cancer types^63^. Thus, we propose that CCM lesion expansion arises from the TRPV2-driven invasive capacity of mutant ECs following lesion onset upon TRPV4-driven endothelial destabilization.

A battery of inhibitors against these MSCs is currently under development. There is no specific inhibitor of PIEZO1 yet, whereas specific and safe TRPV2 and TRPV4 inhibitors already exist. Among them, the candidate TRPV4 inhibitor GSK2798745 has successfully completed Phase 2 safety clinical trials^64^. We showed that it restored the atrioventricular canal in the developing hearts in one-third of CCM2-mutant zebrafish embryos, supporting its capacity to restore cell-cell junction *in vivo*. Tranilast, an anti-allergic medicine used to treat asthma, allergic rhinitis and atopic dermatitis has been widely used as TRPV2 inhibitor^65^. Here, we showed that it prevented the degradation and the invasion of the ECM by CCM2-silenced ECs. Based on these findings, we propose a combined therapeutic strategy to target the two arms of the TRPV2:TRPV4 ratio and prevent CCM onset and progression. Expression levels of these channels also represent promising avenues for identifying biomarkers of CCM disease progression. Such biomarkers are critically needed to guide medical and surgical decision-making and will serve as key surrogate endpoints for assessing treatment efficacy in future therapeutic clinical trials.

This work identifies highly pertinent novel actors at the cell–microenvironment interface. It lays the foundations for a new investigative field focusing on the interplay between aberrant mechanotransduction and CCM genetic and epigenetic defects^66^ to fully understand disease pathogenesis^16^. Beyond CCM, this work redefines the mechanistic landscape of mechanosensitive signaling by uncovering unprecedented functional coupling between PIEZO and TRPV channels. This paradigm shift—from single-channel to multi-channel mechanosensory complexes—has far-reaching implications for the field, offering a new framework to understand how cells decode complex mechanical microenvironments. Our findings may thus inform research in angiogenesis, cancer metastasis, and vascular homeostasis, where PIEZO1, TRPV2, and TRPV4 are individually implicated but whose collective behavior remains largely unexplored.

## Materials and Methods

### Patient Tissue Collection and Preparation

This study protocol was carried out following ethical guidelines established by the Declaration of Helsinki and received formal approval from the local ethics committees at Hannover Medical School (approval no. 6960) and the University of Bonn Medical Center (approval nos. 076/08 and 042/08) in Germany. Written informed consent was obtained from each participant prior to their inclusion in the study. Inclusion criteria required confirmed CCMs based on both neuroradiological imaging and histopathological examination. Intraoperative brain tissue samples from patients undergoing surgical resection of CCMs (n = 42) were collected immediately after excision and preserved on dry ice. As a control group, healthy human brain tissues were obtained from Amsbio. A portion of each sample was snap-frozen and stored at – 80 °C for subsequent gene expression analyses. For histological and immunohistochemical analyses, tissue samples were fixed overnight in 4% buffered formalin solution, processed into paraffin blocks. All patients presented with at least one clinical manifestation related to CCM, including seizures, intracerebral hemorrhage, or focal neurological deficits.

### Total RNA isolation and RNA quality

20 mg of tissue were precisely weighed and transferred into homogenization tubes containing ceramic beads (Precellys system, CK14L; Peqlab) along with 700 µL of RLT lysis buffer from the RNeasy Mini Kit (74104; Qiagen) supplemented with 1% beta-mercaptoethanol. Samples were homogenized utilizing the Precellys 24 Homogenizer (Peqlab) at 6000 rpm with 4 cycles, each lasting 5 seconds. Total RNA extraction was performed following the manufacturer’s instructions for the RNeasy Mini Kit. RNA concentration and purity were assessed using the Nanodrop ND-1000 spectrophotometer. RNA integrity was evaluated using an Agilent Bioanalyzer 2100 equipped with an RNA 6000 Nano Kit (5067-1511; Agilent Technologies).

### Real-time PCR from human tissue samples

Total RNA was reverse transcribed into complementary DNA (cDNA) following the manufacturer’s protocol using the High Capacity cDNA Reverse Transcription Kit (Applied Biosystems, Cat. No. 4368814). Subsequent quantitative gene expression analysis was performed on the StepOne™ Real-Time PCR system, utilizing 15 ng of cDNA template for TRPV4, PIEZO1, and PIEZO2, and 25 ng for TRPV2. Relative gene expression changes between CCM tissues and healthy controls were determined by applying the comparative 2ˉΔΔCt method, normalized against the expression of the housekeeping gene Glyceraldehyde-3-phosphate dehydrogenase (GAPDH). Specific TaqMan® assay IDs used for human tissues were as follows: TRPV2 (Hs00901648_m1), TRPV4 (Hs01099348_m1), PIEZO1 (Hs00207230_m1), PIEZO2 (Hs00926218_m1), CDH5 (Hs00901463_m1) and GAPDH (Hs99999905_m1).

### Immunohistochemistry of human CCM sections

Immunohistochemistry (IHC) was performed on formalin-fixed, paraffin-embedded (FFPE) tissue sections obtained from 30 cerebral cavernous malformation (CCM) specimens surgically resected from patients, alongside 6 commercially sourced normal adult brain samples (Amsbio) with no known cerebral vascular anomalies. Automated immunostaining included positive controls as tonsil tissue sections, known to express these proteins, and negative controls with omission of the primary antibody to confirm specificity and reliability. Primary antibodies utilized were TRPV2 (Abcam, ab272862), TRPV4 (Abcam, ab191580), PIEZO1 (Novus Biologicals, NBP1-78537) all diluted at 1:500. Microscopic imaging was conducted using a Leica microscope equipped with Leica LAS X software at 40x magnification. Five non-overlapping random fields per section were captured for analysis. Semi-quantitative analysis of protein expression was carried out using QuPath software, employing a consistent threshold for DAB chromogen positivity across all samples. Endothelial regions of interest were manually annotated, and DAB-positive cells were identified using a threshold for optical density (OD) set at 0.30. Positive cell counts were expressed as the percentage of DAB-positive endothelial cells relative to the total endothelial cell count within these regions.

### Statistical correlation analysis of lesion size with MSCs expression levels

Clinical data were collected from the cohort of 42 CCM patients. Each patient record included cavernoma volume (in cm³), lesion location (brainstem vs. other), multiplicity (single vs. multiple CCMs), familial vs. sporadic presentation, and patient age. To explore the combined effect of TRPV2 and TRPV4 expression on lesion size, we computed the expression ratio (TRPV2:TRPV4) for each patient. Patients with missing values for either gene, or cavernoma size were excluded. A Pearson correlation analysis was then performed between the TRPV2:TRPV4 ratio and cavernoma size. Additionally, a linear regression model was fitted to visualize the relationship, and the association was plotted with a best-fit line and annotated correlation metrics using ggplot2.

### Stratified MSCs Expression Analysis by Brainstem Status

To investigate the association between brainstem lesion involvement and expression of MSCs, we generated boxplots for PIEZO1, TRPV2, and TRPV4 expression levels stratified by brainstem status (0 = No, 1 = Yes). Given the non-normal distribution of expression data in both groups of unequal sizes, we employed the Wilcoxon rank-sum test for pairwise comparisons.

### Cell culture and siRNA transfections

Pooled Human Umbilical Vein Endothelial Cells (HUVECs) were obtained at passage 0 from Lonza or Pelobiotech. Human Brain Microvascular Endothelial Cells (HBMVECs) were obtained at passage 2 from Pelobiotech. Upon reception, HUVECs and HBMVECs were expanded over 2 or 1 passages respectively in 100µg/ml collagen 1 (from rat tail, BD) coated flasks in complete EGM-2 medium (Lonza) supplemented with 100 U/ml penicillin and 100 μg/ml streptomycin at 37°C in a 5% CO_2_ - 3% O_2_ humidified chamber. HUVEC at passage 3 or HBMVECs at passage 4 were transfected twice at 24 h-intervals with 30 nM siRNA and Lipofectamine RNAi max (Life Technologies, ref. 13778-150) according to the manufacturer’s instructions in OptiMEM at 37°C in a regular 5% CO_2_ humidified chamber. For double transfections, 30 nM of each siRNA duplexes (Dharmacon smartpool ON-TARGET plus Thermo Scientific) was used : Non-targeting siRNA #1 (ref. D-001810-01), CCM2 siRNA (ref. L-014728-01), PIEZO1 siRNA (ref. L-020870-03-0010), PIEZO2 siRNA (ref. L-013925-02-0010), TRPV2 siRNA (ref. L-004194-00-0010) and TRPV4 siRNA (ref. L-004195-00). Inhibitors and agonists were purchased from Bio-Techne/Tocris (GSK1016790A cat. no. 6433, Yoda 1 cat. no. 5586, OB1 cat. no. 6545, Tranilast cat. no. 1098). The vehicle condition corresponds to the dilution factor (or the lowest dilution factor when using multiple drugs in parallel) of DMSO or sterile water in complete medium, depending on the drug resuspension medium.

### Gene expression analysis by RT-qPCR and bulk RNA sequencing

Transfected HUVECs or HBMVECs were cultured for 48 hours after second round of transfection. Cells were harvested and pelleted. Total RNA is extracted from the cells using the NucleoSpin RNA kit (Macherey-Nagel) according to the manufacturer’s instructions. 1µg purified RNA was reverse transcribed into cDNA using the iScript Reverse Transcription Supermix kit (Biorad). Real-time quantitative polymerase chain reaction (qPCR) was performed in 384-well plates using the iTaq Universal SYBR Green Supermix (Biorad), gene-specific primers (see Table), and the CFX-384 instrument (Biorad). For each gene, primers are designed using PrimerBlast or selected from the literature after verifying their specificity on PrimerBlast and ordered from Invitrogen. The specificity (melting curve analysis) and efficiency of the primers are validated prior to experimentation via qPCR on a dilution series of a control RNA sample derived from HUVEC. For the analysis of relative gene expression, each gene was amplified in triplicate for each condition (sample), and three housekeeping genes were also amplified for each condition: GAPDH, RANBP1, and HPRT1. Normalization and quantification of expression were performed using the CFX Manager software, taking into account all housekeeping genes, selecting the siCT condition as a reference, and averaging the technical replicates for each condition.

Gene specific primer sequences used for qPCR:

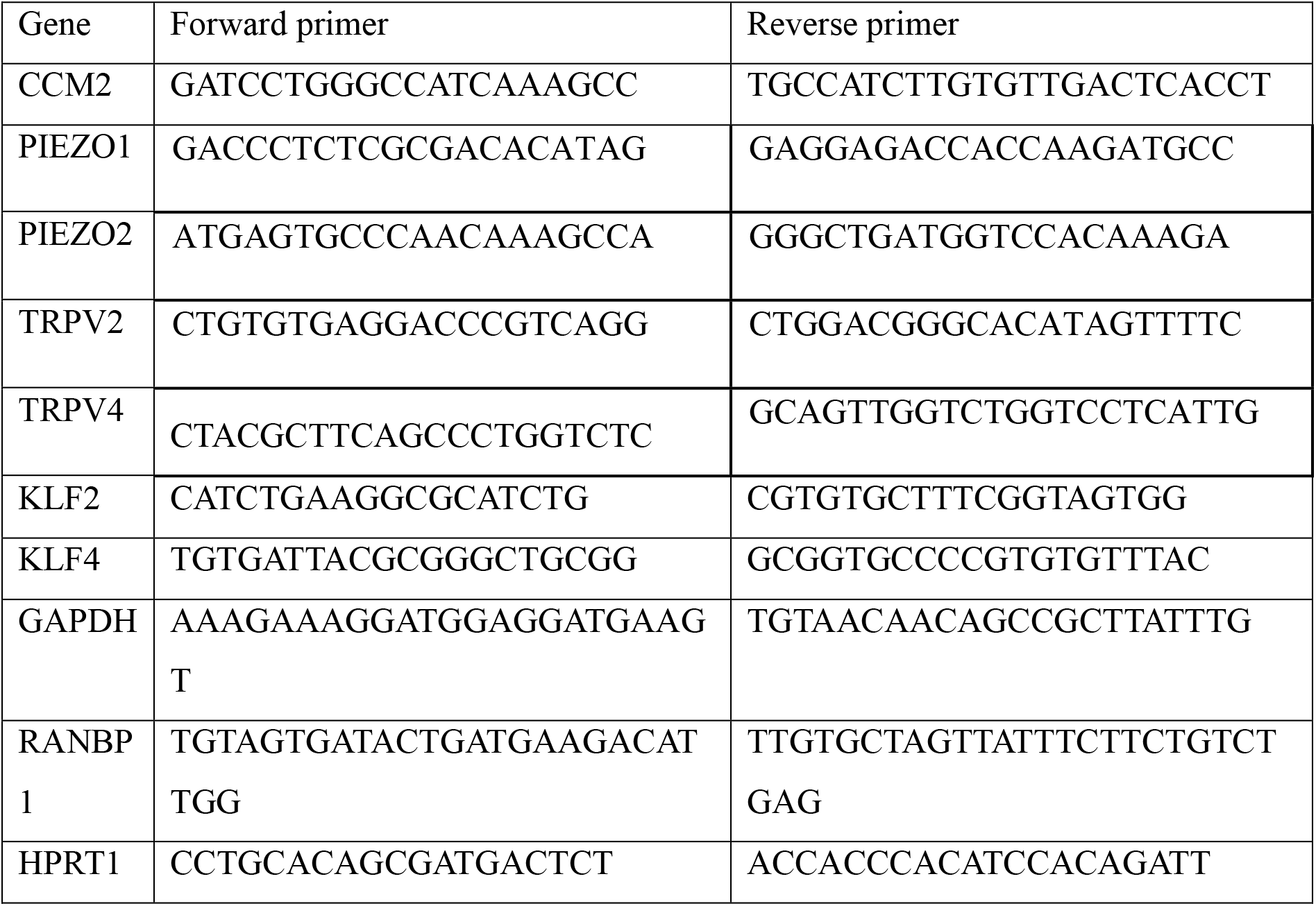

### Bulk RNA sequencing and transcriptomis analysis

Bulk RNA sequencing was performed by BGI Genomics (Shenzhen, China). Total RNA quality and integrity were assessed prior to library preparation. Poly(A)-enriched mRNA libraries were constructed following BGI’s standard protocols, including RNA fragmentation, cDNA synthesis, adapter ligation, and PCR amplification. Libraries were sequenced on a DNBSEQ™ platform using 150-bp paired-end (PE150) reads. Raw sequencing data were subjected to quality control to remove adapter sequences, low-quality reads, and reads containing excessive ambiguous bases. Clean reads were subsequently used for downstream bioinformatic analyses, including alignment to the reference genome, gene expression quantification, and differential expression analysis with DEseq2. Heatmaps using a correlation matrix were obtained with custom R script. DEGs (*P* < 0.05, 5% FDR) rescued by additional silencing of the MSCs were recovered at the union of Venn diagrams of DEGs in siCCM2 vs siCT and DEGs in siCCM2 vs siCCM2 + MSC. Gene Ontology analysis was performed using G-profiler from ELIXIR infrastructure. Gene Set Enrichment Analyses were conducted using GSEA software from Broad Institute, UC San Diego.

### Western Blots

Transfected cells were harvested 48h after second round of transfection, lysed in RIPA buffer, and sonicated. Protein concentration was determined using a BSA standard curve (Micro BSA kit). Equal amounts of protein were mixed with Laemmli buffer, heated, loaded, and separated on an SDS-PAGE gel, then transferred to PVDF membranes. The following antibodies were used: TRPV2 clone 2D6 (diluted 1:1000, Sigma, MABN2426), TRPV4 clone 1B2.6 (diluted 1:500, Sigma, MABS466), PIEZO1 (diluted 1:500, Novus NBP2-75617), KLF4 (Invitrogen, MA5-35030), GAPDH (Cell Signaling, cat. no. 2118T). Immunodetection was performed using an HRP-conjugated secondary antibody. Peroxidase activity was visualized with Clarity ECL substrate (Bio-Rad) using a Fusion FX imaging system (Vilber). Densitometric quantification of the bands was performed using the Evolution Capture software. Protein levels were normalized relative to the GAPDH level.

### Calcium imaging experiments

Transfected HUVECs were seeded 24 h after the second round of transfection in collagen-coated black 96-well clear glass bottom plates (4×10^4^ cells/well) in complete EBM-2 medium (Lonza) supplemented with 100 U/ml penicillin and 100 μg/ml streptomycin at 37°C in a 5% CO2 - 3% O2 humidified chamber overnight. Cells were loaded with 2,5µM Fura-2-AM probe (F1201, Fisher Scientific) diluted in complete media for 45 minutes at 37°C. After loading, cells were washed twice in HBSS (142.6 mM NaCl, 5.6 mM KCl, 5.6 mM glucose, 10 mM HEPES, 0.34 mM Na_2_PO_4_, 4.2 mM NaHCO_3_, 0.44 mM KHPO_4_, pH 7.4) supplemented with 1.8mM CaCl_2_ and 0,8mM MgCl_2_. Fluorescence emission at 510nm was acquired on the FlexStation3 (Molecular Devices) every 2s following FURA-2 excitation at 340 and 380 nm and analyzed with the SoftMax Pro software (Molecular Devices). A baseline was established for 1 min then 2.5 μM Yoda1 (PIEZO agonist) or 25nM GSK1016790A (GSK101; TRPV4 agonist) was added. After 2min, 1μM of the ionophoric drug ionomycin was added to force Ca^2+^ entry and check cell survival.

### Immunofluorescence staining

Transfected HUVECs were seeded at 1.5×10^6^ cells in 24-well plates on coverslips coated with 10 μg/ml fibronectin (from human plasma, Sigma Aldrich) for 48h in complete EBM-2 medium. Cells were fixed with 4% PFA, permeabilized with 0.2% Triton X-100, and incubated with antibodies against anti-βcatenin (mouse monoclonal antibody C7082 Sigma Aldrich, 1/500) or vinculin (mouse monoclonal antibody V4505 Sigma Aldrich, 1/200). After rinsing, coverslips were incubated in Goat anti-Mouse IgG (H+L) highly cross-adsorbed secondary antibody, Alexa Fluor conjugated AF 488 (Invitrogen, 1/1000) and Alexafluor647 phalloidin (Invitrogen A2287, 1/2000). The coverslips were mounted in Mowiol/DAPI solution and imaged on a Zeiss Axio Imager epifluorescence microscope equipped with a Hamamatsu monochrome camera, using a 20X objective (1037 x 1387 pixels; pixel size 0.317 µm) or 63X objective under oil immersion (pixel size 0.1031). Ten fields were randomly acquired per condition.

### Cell-cell junction thickness quantification

To quantify the width of cell-cell junctions, the β-catenin signal from the 20X images was segmented using Ilastik pixel classification to generate a binary junction mask. Junction width was then extracted in Fiji by multiplying the Euclidean distance map (EDM) of the binary mask by its skeleton, so that each skeleton pixel carries the local half-width of the junction it belongs to. Half-width values were converted to micrometers using the pixel size, pooled over all skeleton pixels of all fields of a given condition, and conditions were compared through their cumulative distributions of junction half-width.

### Cortical/cytosolic actin ratio quantification

The cortical-to-cytosolic F-actin ratio was quantified on the same β-catenin/phalloidin image sets used for junction width analysis. Individual cells were segmented on the β-catenin channel using Cellpose with the cyto3 model, with automatic diameter estimation kept constant across all images of a given replicate; cells whose mask touched the image border were excluded so that only fully imaged cells were retained. The resulting single-cell ROIs were transferred onto the corresponding TRITC-phalloidin channel and the total F-actin raw integrated density was measured within each ROI. Each ROI was then eroded by 20 pixels (6.3 µm) to define the cytosolic compartment, and the cortical compartment was defined as the resulting 6.3 µm-wide peripheral band. Cortical F-actin was calculated as the difference between the total and the cytosolic raw integrated densities, and the cortical-to-cytosolic ratio was computed for each individual cell. On average 60 cells were analysed per field.

### Permeability assay

Twenty-four-well plates were coated with biotinylated gelatin diluted in sodium bicarbonate buffer to a final concentration of 25 mg/mL and incubated for 1 h at 37°C. HUVECs were then seeded at a density of 150,000 cells per well and cultured for 48 h to allow the establishment of cell-cell junctions. To assess endothelial permeability, cells were incubated for 3 min with serum-free Opti-MEM containing FITC-conjugated streptavidin at a final concentration of 25 µg/mL. The medium was then aspirated and cells were immediately fixed with 4% paraformaldehyde (PFA) for 15 min. After three washes, cells were permeabilized with 0.2% Triton X-100 for 10 min and subsequently incubated with TRITC-phalloidin at 1:25000 in BSA-containing buffer for 30 min, then a Mowiol/DAPI solution was added to the wells. Ten-fourteen fields were randomly acquired per condition using an inverted microscope at 40× magnification. Image segmentation, processing and quantification were carried out in Fiji/ImageJ. A binary mask of the streptavidin fluorescence was generated, and the positive area was quantified for each image and expressed as a percentage of the total image area.

### Focal adhesion quantification

To calculate the focal adhesion count and size, the vinculin signal from 63X images were first automatically segmented using Ilastik followed by analysis through an in-house developed MATLAB code. The segmented images were first cleaned up by removing any regions with a surface area smaller than 0.0744 µm² as well as any regions with a length along the long axis less than 0.5155 µm. Next, the number of focal adhesions was calculated by counting the number of remaining objects per field of view (FOV). The average focal adhesion size was obtained by calculating the mean size of all the objects inside a FOV, multiplied by the pixel surface size in µm².

### Gelation degradation assay

Coverslips in 24 well-plates were coated with gelatin-Alexa488 dye as previously described ^67^. Transfected HUVECs were seeded 24h after the second siRNA transfection at 3×10^4^/well in OptiMEM medium in presence or absence of the calcium channels inhibitors overnight at 37°C in a 5% CO_2_ incubator. Cells were fixed with 4% PFA, washed with PBS twice. The coverslips were mounted in Mowiol/DAPI solution and imaged on an on a Zeiss Axio Imager epifluorescence microscope equipped with a Hamamatsu monochrome camera using a 63X objective under oil immersion (pixel size 0.1031) Ten fields were randomly acquired per condition. Images were converted to binary images using B&W thresholding on ImageJ. Images were automatically segmented using Ilastik pixel classification to generate a binary degradation mask. The total area of the black zones corresponding to the total area of degradation of the fluorescent gelatin was measured and normalized to the number of nuclei.

### Angiogenic invasion assay

24 hours after the second round of siRNA transfection, PEG-based 3D invasion samples were prepared following our previously published protocol^30^. For samples that were fixed directly after the cell invasion, a 1.5% PEG hydrogel containing 80-120nM sphingosine 1-phosphate (S1P, Sigma-Aldrich, S9666) and lysine-RGD (pepMic) without microspheres was used (E-modulus of 584Pa). For TFM experiments, dark red fluorescent 200nm microspheres (ThermoFisher Scientific, F8807) were included in a 1.2% PEG hydrogel (E-modulus of 330Pa). E-moduli were derived from bulk shear rheological measurements on cylindrical gel samples (7mm diameter, 1mm height, n≥3 for both gel formulations) that were allowed to swell overnight in 1× PBS. Rheological measurements were performed at 37 °C using an MCR 702 rotational rheometer (Anton Paar) equipped with a P-PTD200/80/I Peltier bottom plate. Samples were placed between roughened aluminum surfaces and compressed to ∼80% of their original height (0.8-mm gap) to minimize slippage^68^. Shear storage moduli (G’) were obtained for 3min time sweeps at 1% engineering shear strain (within the linear range, as verified by strain sweeps) and 1 rad s⁻¹ and converted to Young’s moduli by assuming a Poisson’s ratio of 0.3.

For invasion distance quantification and immunostainings, 40.000 cells were seeded on the PEG gel. For TFM experiments, the cells were first stained with 2µM green CellTracker (Fisher Scientific, Invitrogen C7025) prior to their seeding on the gel. After cell seeding, the imaging plates were moved to the incubator and stored vertically for one hour to allow evenly distributed cell attachment to the gel surface. After this hour, the plates were flipped horizontally, 1mL medium was added to each sample and the samples were incubated at 37°C for 16 to 18 hours in a 5% CO2 - 3% O2 humidified chamber.

### Traction Force Microscopy imaging

16 to 18 hours after cell seeding on the PEG gels, TFM-samples were moved from the incubator to a Leica SP8 confocal microscope equipped with stabilized incubation (37°C, 5% CO_2_) and stage control. Images were made with a 20x 0.4 NA dry objective at 2X zoom with a 512×512 resolution. Voxel-size was 0.568 x 0.568 x 1 µm^3^.

For each sample, ten positions of invading cell sprouts were selected based on sprout appearance (healthy-looking sprouts) and amount of invasion in the selected region (preferably isolated sprouts). Image stacks with 1µm z-step were created to ensure enough beads were captured to detect all cell-induced gel-deformations. ‘Stressed’ images of the active cells were captured per sample, after which immediately Cytochalasin D (CytoD, Avantor, 22144-77-0) in a final concentration of 4µM was added to the imaged well using a syringe. Images for all six cell conditions were captured. One hour after first CytoD addition, the imaging sequence was restarted for all samples to capture the ‘relaxed’ images, ensuring exposure to CytoD and relaxation of all samples for at least 50 minutes.

### Traction Force Microscopy processing

TFM analysis was preformed using our in-house developed TFMLAB toolbox in MATLAB and as described earlier^30,69^. First, the bead and cell images were filtered and enhanced and cells were segmented to create a cell mask. Next, shift correction was performed using the Gradient Based (fast first order Taylor) algorithm. After this, the displacement field was calculated using free form deformation image registration. These displacement fields were exported for the full field of view of the images and visualized in ParaView in order to make first assessment of the quality of the data, looking out for outliers and irregular fields. For trustworthy images, crops were then made around sprout tips of interest. For these regions, tractions were calculated using a previously established and validated physics-based nonlinear inverse method (PBNIM) implemented into a finite element framework^70^, and taking into account the mechanical characteristics of the hydrogel (E-modulus 330Pa, Poisson’s ratio 0.3). To account for outliers, 99^th^ percentile displacements and tractions are reported, as well as the strain energy of the sprout tips. Sprouts, displacement fields and traction fields were visualized in ParaView.

### Immunofluorescence staining of 3D invasion

Samples that were used to measure the invasion distance were stained overnight at 4°C with 1:1000 phalloidin-Atto 647N (Sigma-Aldrich, 65906) and with 1:2000 DAPI (ThermoFisher Scientific, D1306) for 5 to 10 minutes at room temperature. For immunofluorescence staining of pMLC, the adhesive chambers were carefully removed from the glass bottom dish and flipped upside down. Samples were first permeabilized with 0.3% Triton X-100 (Sigma-Aldrich, T8787) in PBS for 1 hour at room temperature. Next the samples were incubated in permeabilizing blocking solution (1% BSA, 10% goat serum, 1% Triton X-100, PBS) for at least one hour. Primary antibody 1:50 Phospho-Myosin Light Chain 2 (Ser19) Antibody (rabbit, Cell Signaling Technology, 3671) and secondary antibody 1:500 Goat anti-Rabbit IgG (H+L) Highly Cross-Adsorbed Secondary Antibody, Alexa Fluor 546 (Invitrogen, A11035) were incubated overnight at 4°C. After each incubation, the samples were washed with 0.05% TWEEN20 in PBS every 2 hours for 8 hours. The chambers were then carefully flipped again on a cover glass-bottom well plate and stored in PBS at 4°C until imaging.

### Invasion distance quantification

For the invasion analysis, overview images of the entire cell layer of fixed, phalloidin- and DAPI-stained samples were acquired through confocal microscopy. The images were made on a Leica SP8 confocal microscope using a 10x 0.4 NA dry objective with a 512×512 resolution. Voxel-size was 2.275 x 2.275 x 1 µm^3^. 100µm thick stacks were acquired in order to capture sufficient sprouts for all cell conditions.

To quantify the invasion distance, maximum projections of the cell layers were created in imageJ. On this image the distance between the cell layer and the farthest invaded tip of cell sprouts or individually invaded cells was measured perpendicular to the cell layer. This was done over the complete length of the cell layer, always selecting the farthest invaded cell in the selected area. All invasion distances were averaged per monolayer and normalized to the control condition per biological replicate to account for differences in fixation timing and S1P variability.

### Actin and filopodia quantification

To quantify the actin and filopodia, close-up images of cell sprouts were acquired of phalloidin- and DAPI-stained samples. Images were made on a Leica SP8 confocal microscope using a 25x 0.95 NA water objective with a 1024×1024 resolution. The voxel-size was 0.151 x 0.151 x 1 µm^3^. Per sample per trial, 5 to 10 sprouts were selected, ensuring the completed sprout was captured in the stack.

Image processing for actin and filopodia was done in MATLAB. First, a cell mask of the sprout of interest was made by manually creating a crop on a binarized maximum intensity projection of the image. For the actin intensity quantification, the 3D image was first filtered with a median filter (medfilt3). Next, the values of the image outside of the cell mask in the 3D image were set to zero. The normalized total actin intensity was calculated by summing all intensity values in the 3D image and dividing by the number of nuclei in the cell sprout. Results were normalized to the control per trial. For the number of filopodia in the sprout tips, an opening operation using a disk-shaped element with a 7-pixel radius was applied to the 2D binary cell mask. The resulting image was subtracted from the 2D cell mask and elements smaller than 50 pixels were removed. The remaining separate structures in this image were counted and divided by the number of nuclei in the cell sprout.

### Angiogenic sprout immunostaining quantification

For the quantification of pMLC, close-up images of cell sprouts were acquired on a Leica SP8 confocal microscope using a 25x 0.95 NA water objective. Within each biological replicate the same resolution was used, either 1024×1024 (voxel-size of 0.151 x 0.151 x 1 µm^3^) or 512×512 (voxel-size of 0.303 x 0.303 x 1 µm^3^). 5 to 10 sprouts were selected per condition, ensuring the complete sprout was captured in the stack.

Image processing of the immunofluorescence images was done in MATLAB. A cell mask of the sprout of interest was manually selected on a binarized maximum projection of the image. Next, the 3D image was filtered with a median filter (medfilt3). All pixels outside the cell mask in the 3D image were set to zero, after which all pixels were summed and dividing by the number of nuclei. This total intensity per cell was normalized to the control per trial.

### Zebrafish handling, oocyte injections, and pharmacological treatments

Zebrafish handling was performed in compliance with German and Brandenburg state law, carefully monitored by the local authority for animal protection (LAVG). *ccm2^m201 (34)^* and *Tg(kdrl:EGFP)^s843 (71)^* zebrafish lines were used. Antisense oligonucleotide morpholino (MO) *trpv4* (5’-GTTACAAAGAAAAAGAGTCCAGAAC-3’; Gene Tools, LLC, Eugene, Oregon;^55^) was injected with 1.8 ng per embryo at the one-cell stage and embryos were then incubated until 54-56 hpf at 28 °C in E3 medium. For pharmacological treatment of zebrafish embryos, GSK2798745 (HY-19765, MCE) was used. The compound was dissolved in DMSO to produce 10 mM of stock solution. At the 16 somite-stage, approximately 30 dechorionated embryos were transferred per well into 6-well plates and supplied with 100 μM GSK2798745 in 4 ml E3 medium. Control embryos were treated with 1% DMSO in E3 medium. Embryos were then incubated until 54-56 hpf at 28.5°C.

### Zebrafish Immunohistochemistry and imaging

Zebrafish whole-mount immunohistochemistry was performed as previously described^72^. Embryos were fixed with 4% PFA overnight and permeabilized with PBST with 0.5% Triton X-100 for 1 hour and subsequently incubated with primary antibody mouse anti-ALCAM (1:200, DSHB AB_531904) diluted in PBST with 0.2% Triton X-100, 1% BSA, and 5% NGS. Secondary antibodies were Alexa Fluor 633-conjugated goat anti-rabbit (1:200, Invitrogen A21070. Hearts were dissected, flat-mounted in SlowFade™ Glass (Invitrogen, S36964), and imaged using LSM 880 confocal microscopes (Zeiss). Image processing was performed with Imaris v9.1.5 (Bitplane, UK) and Fiji software.

### Statistical tests

All data are expressed as mean ± standard error of the mean (SEM). Statistical significance was assessed by either performing a t-test for comparing 2 conditions or for more than 2 conditions by multiple comparison adhoc tests post-ANOVA to compare with siCT and siCCM2; a 0.5 alpha level was used for all comparisons. P<0.05 was considered to be significant. *P<0.05, **P<0.05, ***P<0.005. n represents biological replicates. Prism software was used to conduct the statistical analysis of all data.

## Acknowledgements

We thank the MicroCell microscopy facility in IAB Grenoble and the members of the Faurobert group for their support. This study was supported by ERANET NEURON JTC 2022 call-funded by ANR, FWO and BMBFR (E.F., H.V.O., S.K.); ANR-22-CE14-0062-02 (E.F.); INCA PLBIO21-215 (E.F.); Research Foundation Flanders (FWO) G0C2422N (H.V.O., E.F., S.R.); Ambition Internationale, Région AURA (E.F., H.V.O., S.K., S.R.); Région Nouvelle Aquitaine 16/RALPC-P-R-23 (A.P.). H.V.O. was supported by KU Leuven internal funding BOF C14/24/112 and iBOF/21/083 C. C.P was supported by a CIFRE PhD fellowship from ANRT and S.V. by KU Leuven Facultaire Luik Onderzoeksfonds (FLOF) and FWO G0C2422N. E.F. is grateful to the French CCM Patient organization ACC and Espoir Isère contre le Cancer for their support.

## Author contributions

J.D.J. and A.S. designed 3D in vitro experiments, which were performed and analyzed by J.D.J. C.P., T.M.R., E.R. and C.P.V. performed and analyzed 2D in vitro experiments. A.A. performed and analyzed experiments on clinical samples. I.P., H.B. and A.S. provided patient clinical data and CCM tissue. N. K., F.F. and C.H. brought support on automated IHC staining and scoring. A.O. conducted statistical and correlation analyses on the human dataset. S.A.S. provided the zebrafish ccm model and C.R. performed in vivo zebrafish experiments. E.B. and C.O. performed molecular biology experiments. F.S., A.P.B. and S.V. developed image analysis tools. S.L.P. and A.P. supervised calcium imaging experiments and analyses. A.P., O.D., E.P., S.R. and S.A.S. contributed to scientific discussions. E.F., S.K. and H.V.O. conceptualized and supervised the study, designed and interpreted experiments. E.F. wrote the manuscript which has been revised by all the authors.

