## Supplementary material for "Endothelial gain of PIEZO/TRPV mechanosensors drives cavernous malformations": sup figures

a

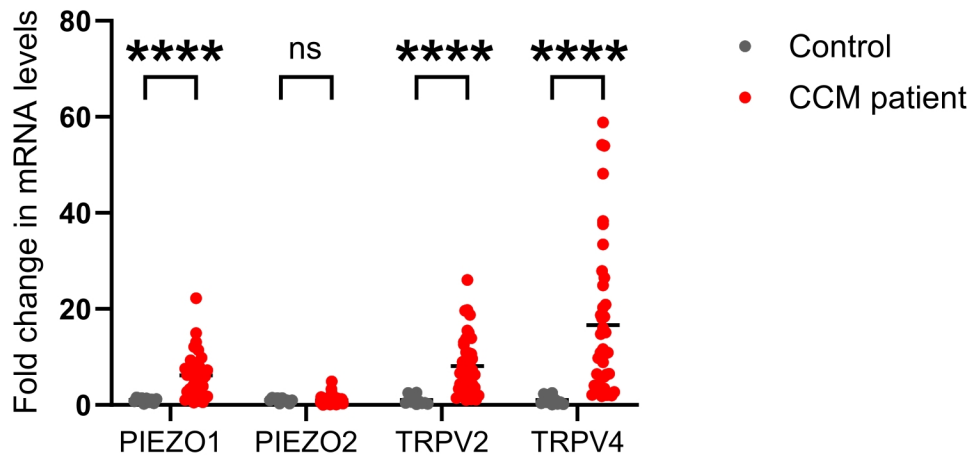

b

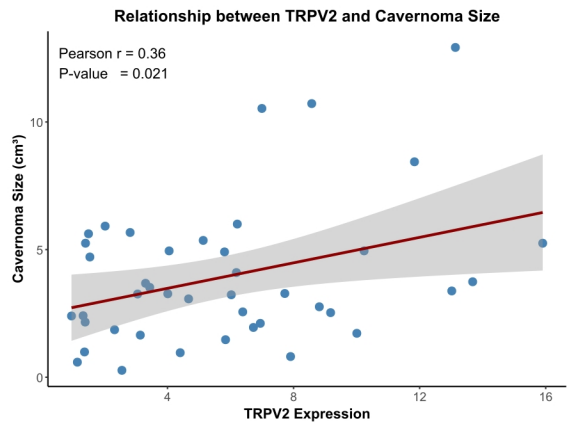

**Figure S1: (a)** : Fold change of PIEZO1, PIEZO2, TRPV2 and TRPV4 mRNA levels in CCM lesions ($n = 42$, red) relative to control brain tissue ($n = 10$, gray), measured by qRT -PCR and normalized to GADPH. Each dot represents an individual patient; bars indicate means. Mann Whitney test; $*p < 0.05$, $****p < 0.0001$. Exact $p$ -values are: TRPV2: $<0.000001$; TRPV4: $<0.000001$; PIEZO1: $0.000005$; PIEZO2: $0.199549$ between controls and CCM patients. **(b)** : Positive correlation between TRPV2 mRNA expression and lesion volume (cm<sup>3</sup>) in patients with CCM lesions ($n = 42$). Pearson $r = 0.36$, $p = 0.021$; red line, linear regression; 95% confidence interval.

a

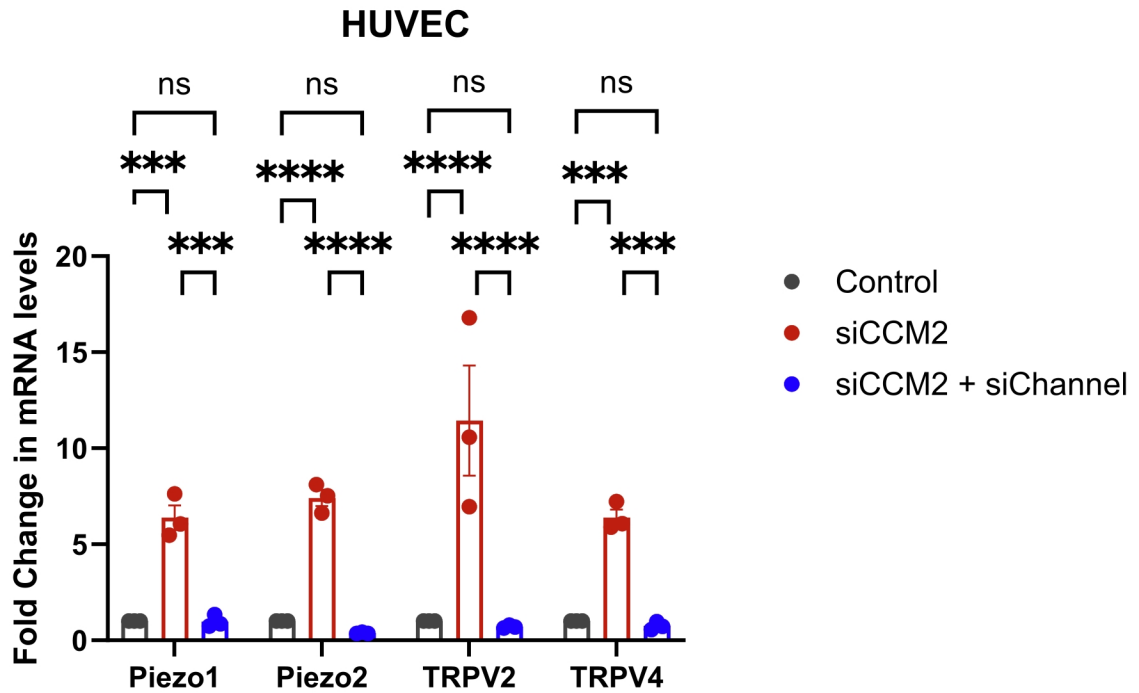

b

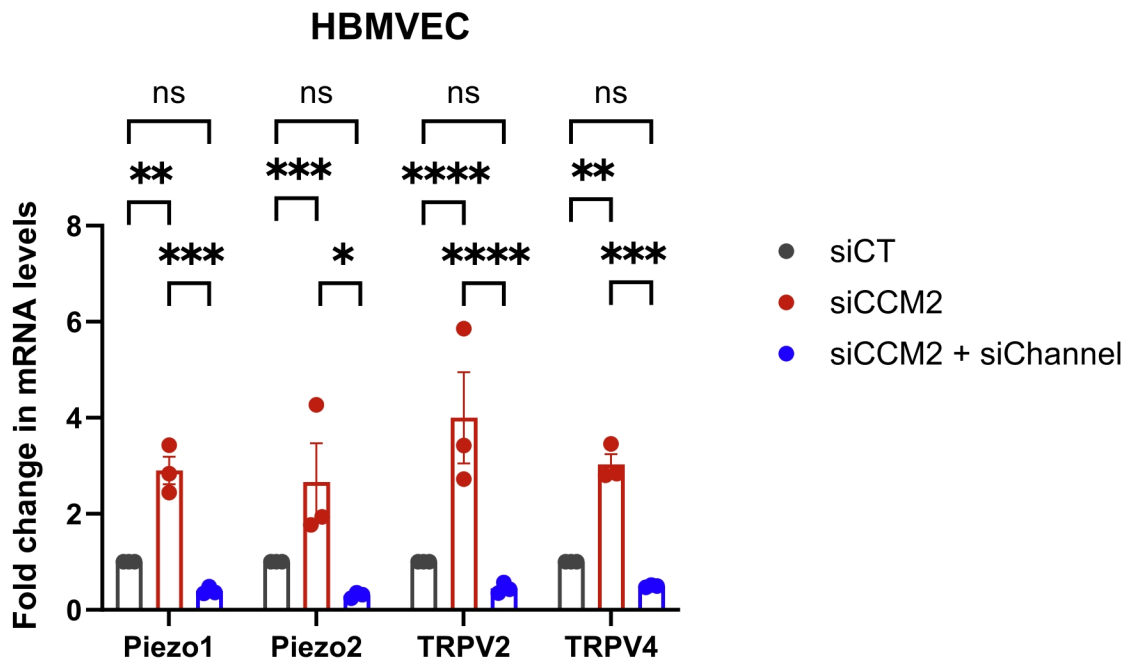

**Figure S2:** Efficiency of silencing of the corresponding channel in doubly silenced HUVECs (a) and HBMVECs (b). Dots are the mean of three technical replicates. ($n=3$ independent experiments). Bars indicate mean ± SEM. Two-way A NOVA with Šídák's correction. \*$p < 0.05$, \*\*\*\*$p < 0.0001$. Exact p-values between siCCM2 and siCCM2+siChannel: in (a) are PIEZO1: 0.0005; PIEZO2 and TRPV2: $<0.0001$; TRPV4: 0.0003. in (b) are PIEZO1 and TRPV4: 0.0002; PIEZO2: 0.0005; TRPV2: $<0.0001$.

A

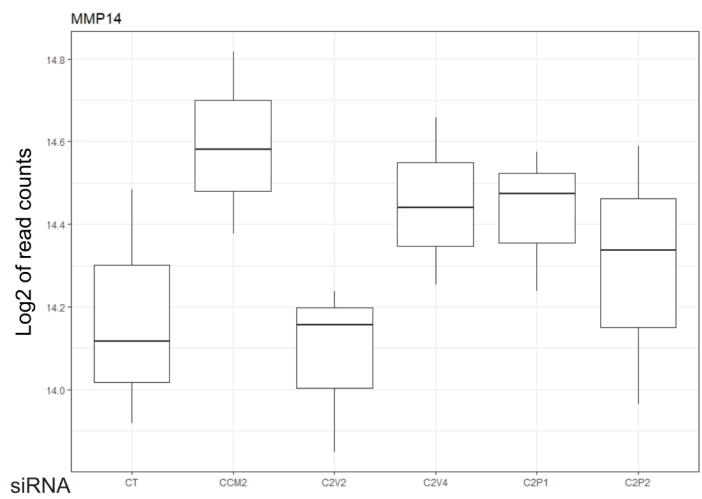

B

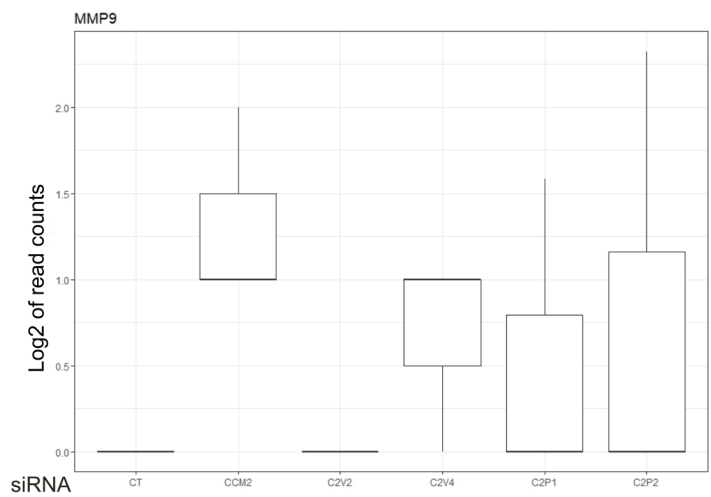

**Figure S3:** Expression of MMP14 (a) and MMP9 (b) in HUVECs in the different silencing conditions as measured by RNA sequencing.

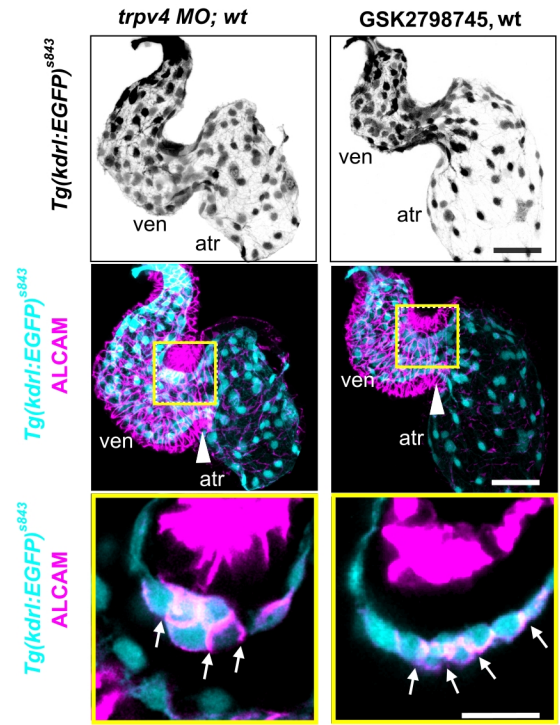

**Figure S4: Effect of TRPV4 morpholino and GSK2798745 on WT zebrafish embryos.** The morphology of wild-type endocardium at 56 hpf, marked by *Tg(kdrl:EGFP)<sup>s843</sup>* expression is not affected by TRPV4 morpholino injection or treatment with GSK2798745.

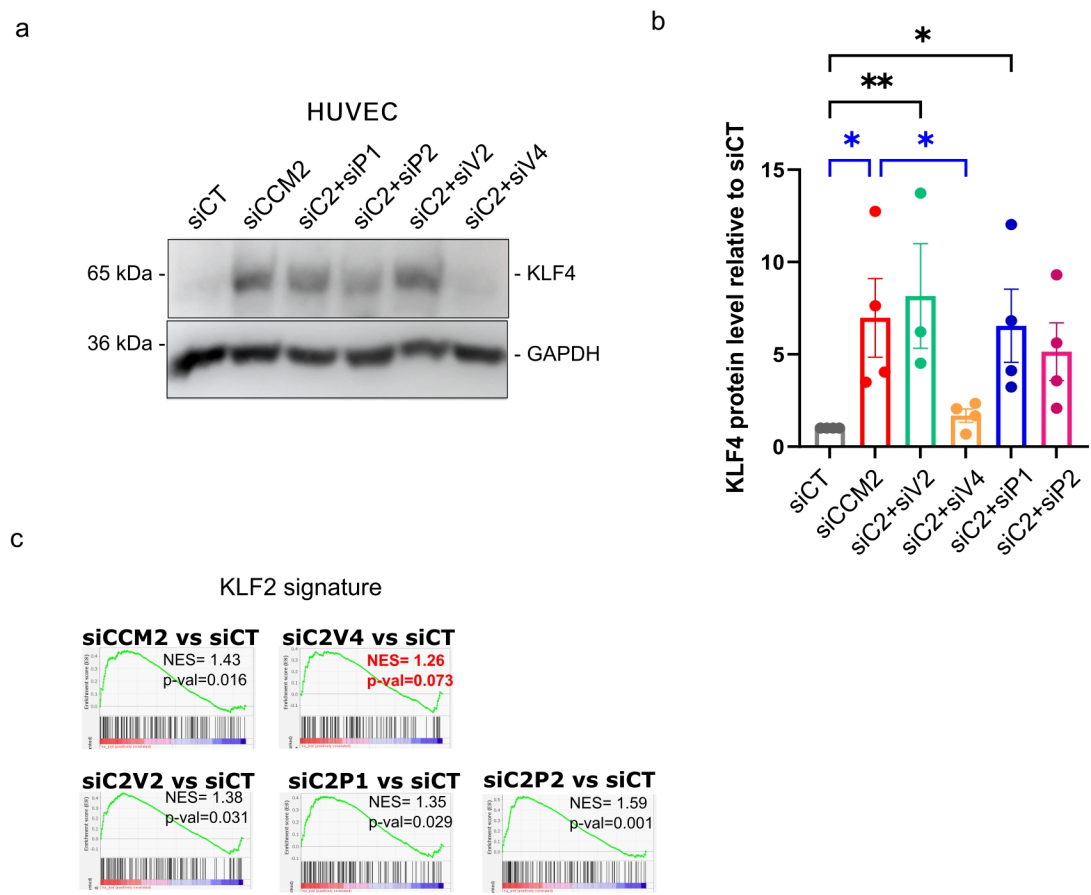

**Figure S5: (a):** Representative images of western blot of KLF4 in silenced HUVEC. GAPDH is shown as a loading control. **(b):** KLF4 normalized protein levels detected by western blot in HUVEC. n=4 independent experiments. Bars indicate mean ± SEM. Two-way ANOVA with Tukey's correction. \*p < 0.05, \*\*\*\*p < 0.0001. Exact p -values are 0.0219 (siCT vs siCCM2); 0,0038 (siCT vs siC2+siV2); 0.0357 (siCT vs siC2+siP1); 0.0477 (siCCM2 vs siC2+ siV4). **(c):** Gene set enrichment analysis of KLF2 signature (from <sup>73</sup>) in the transcriptome of silenced HUVECs using GSEA software from Broad Institute UC San Diego. NES: Normalized enrichment score.
